# Single-cell analysis of signalling and transcriptional responses to type I interferons

**DOI:** 10.1101/2023.07.03.547491

**Authors:** Rachel E. Rigby, Kevin Rue-Albrecht, David Sims, Jan Rehwinkel

## Abstract

Type I interferons (IFNs) play crucial roles in antiviral defence, autoinflammation and cancer immunity. The human genome encodes 17 different type I IFNs that all signal through the same receptor. Non-redundant functions have been reported for some type I IFNs. However, whether different type I IFNs induce different responses remains largely unknown. We stimulated human peripheral blood mononuclear cells (PBMCs) with recombinant type I IFNs to address this question in multiple types of primary cells. We analysed signalling responses by mass cytometry and changes in gene expression by bulk and single-cell RNA sequencing. We found cell-type specific changes in the phosphorylation of STAT transcription factors and in the gene sets induced and repressed upon type I IFN exposure. We further report that the magnitude of these responses varied between different type I IFNs, whilst qualitatively different responses to type I IFN subtypes were not apparent. Taken together, we provide a rich resource mapping signalling responses and IFN-regulated genes in immune cells.

**Highlights:**

- Mass cytometry and scRNAseq analysis of human PBMCs stimulated with type I IFNs
- Cell type-specific phosphorylation of STAT proteins and induction of ISGs
- Different type I IFNs induce qualitatively similar responses that vary in magnitude
- Identification of ten core ISGs, up-regulated by all cell types in response to all type I IFNs

**In brief:** Rigby *et al.* provide a single-cell map of signalling and transcriptomic responses to type I IFNs in *ex vivo* stimulated human PBMCs. Different cell types responded in unique ways but differences between different type I IFNs were only quantitative. These rich datasets are available via an easy-to-use interactive interface (https://rehwinkellab.shinyapps.io/ifnresource/).

**Graphical abstract:** 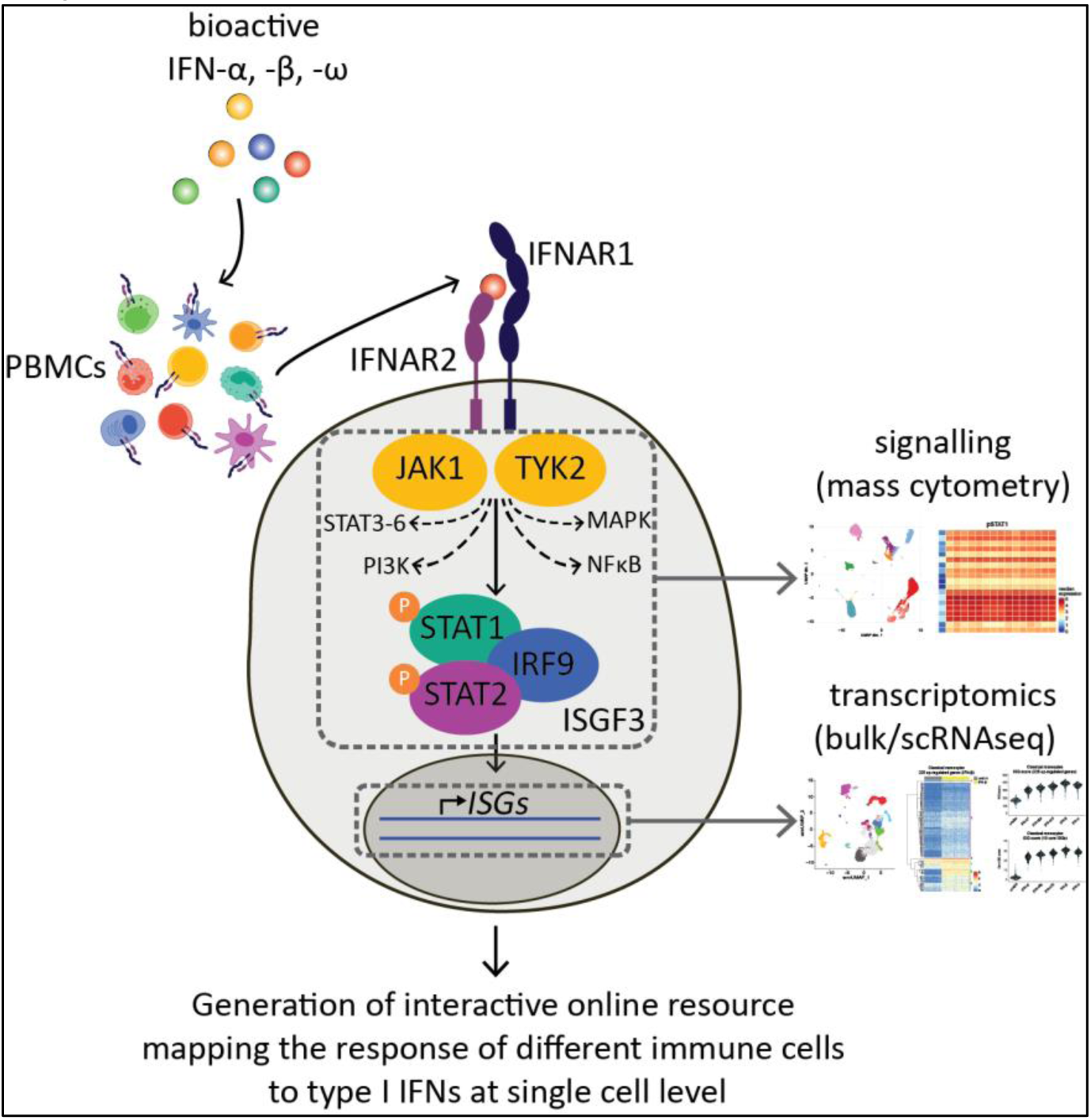

## Introduction

Type I IFNs are a family of cytokines essential to antiviral immunity.^1^ Typically, type I IFNs are produced transiently following an infection. However, in some autoimmune and autoinflammatory conditions, type I IFNs are secreted chronically and initiate and/or exacerbate disease.^2^ Furthermore, type I IFNs impact on bacterial infections and cancer, with both beneficial and detrimental effects.^1,3^ Therefore, type I IFNs have wide-ranging effects in the immune system and in diseases.

Type I IFNs were identified in the 1950s for their ability to “interfere” with virus infection.^4,5^ Much research has been done since on type I IFNs and led to the concept of the type I IFN system (Figure 1a).^6^ This host defence response is triggered when cells autonomously detect virus infection. Nucleic acids are often a molecular signature of viral infection. Nucleic acid sensors recognise viral DNA or RNA, or disturbances to the homeostasis of cellular nucleic acids, such as mis-localisation.^7^ These sensors then initiate signalling cascades resulting in the expression and secretion of type I IFNs. Like in other species, there are many different type I IFNs in humans: 13 IFN-α sub-types, IFN-β, -ω, -ε and -κ. Type I IFNs engage IFNAR, a receptor expressed on the surface of all nucleated cells.^8^ IFNAR is composed of two subunits, IFNAR1 and IFNAR2, which are associated at their cytoplasmic tails with the kinases TYK2 and JAK1, respectively. The canonical signalling pathway downstream of IFNAR involves the formation the ISGF3 complex by phosphorylated STAT1 and STAT2 together with IRF9. ISGF3 binds to IFN-stimulated response elements (ISREs) in the promoters of type I IFN-responsive genes and thereby drives the expression of IFN-stimulated genes (ISGs). Non-canonical signalling pathways downstream of IFNAR can involve the phosphorylation of other members of the STAT transcription factor family (STAT3-6), members of the MAP kinase family, components of the PI3K/mTOR pathway, or NF-κB (Figure 1a).^8–14^ Canonical and non-canonical IFNAR signalling profoundly affect the transcriptome.^15^ Initial conservative estimates suggested that ∼400 genes are induced by type I IFN^16^ and more recent experiments in human fibroblasts found that ∼10% of transcripts are controlled by type I IFN.^17^ Moreover, type I IFNs also downregulate mRNA levels of some genes.^17,18^ Some ISGs encode virus restriction factors, cellular proteins that block infection, whilst others encode proteins involved in virus detection or in cellular and adaptive immune responses.^19,20^ Collectively, the genes regulated by type I IFNs thus complete the “loop” of the type I IFN system (Figure 1a).

**Figure 1.**
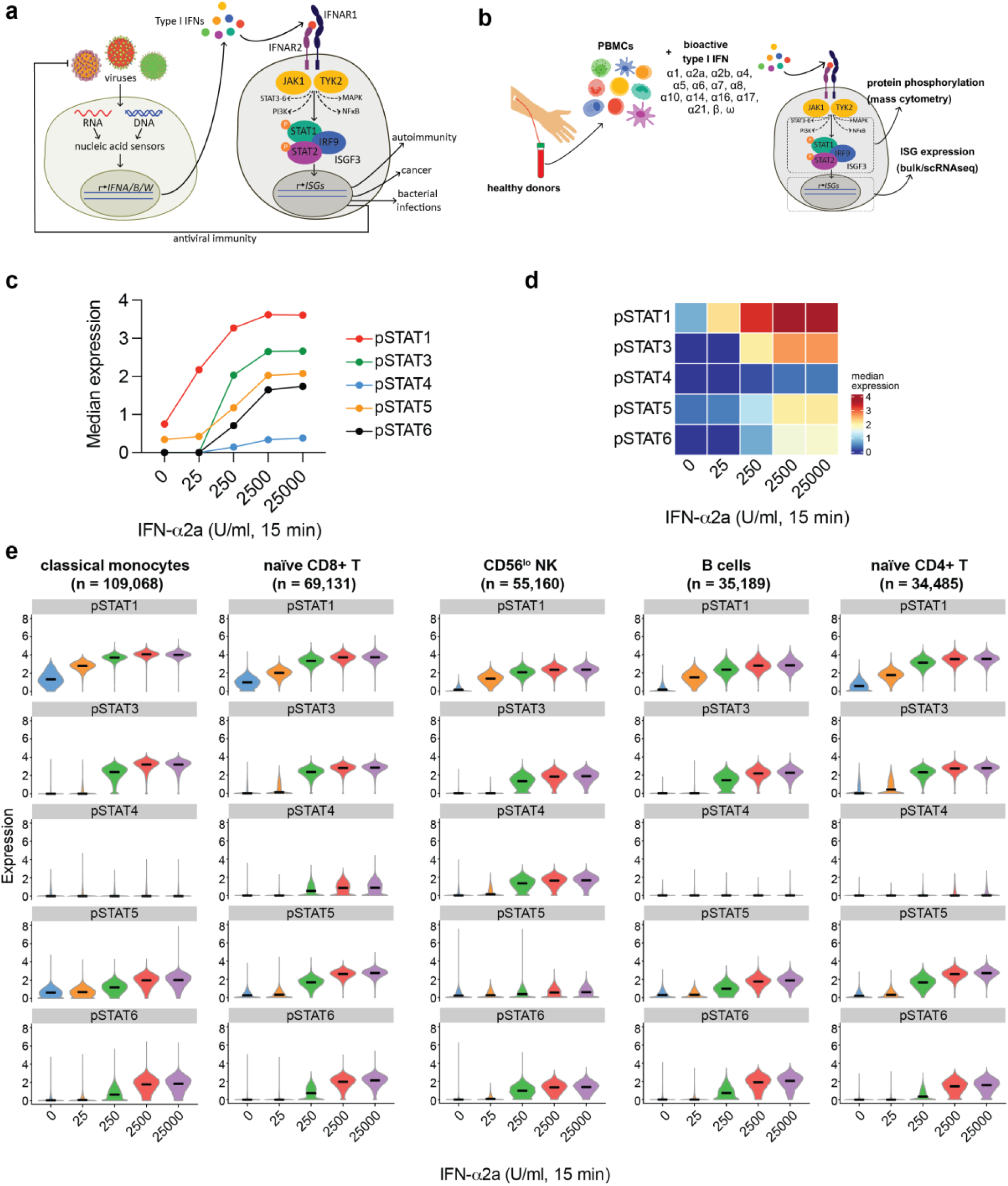
Phosphorylation of STAT proteins in response to IFN-α2a. (a) Overview of the type I IFN system. (b) Schematic representation of the mass cytometry workflow to quantify protein phosphorylation in response to type I IFN-treatment. (c) Median expression of pSTAT1, pSTAT3, pSTAT4, pSTAT5 and pSTAT6 in PBMCs treated with increasing concentrations of IFN-α2a for 15 minutes. (d) Depiction of the data shown in (c) as a heatmap.(e) Violin plots showing expression of each pSTAT in the indicated cell types in response to increasing concentrations of IFN-α2a. The number of cells analysed is shown in parentheses. Violin plots for all cell types are shown in Figure S1c. Data are from one experiment with one donor. See also Figures S1 and S2 and Supplementary Tables 1 and 2.

All type I IFNs bind to and signal through IFNAR, raising the question whether these cytokines have redundant functions. Multiple explanations likely exist for the large number of type I IFNs. These include tissue-specific functions. Indeed, IFN-ε is expressed in the female reproductive tract and may play a role in cancer immunosurveillance.^21,22^ IFN-κ is expressed in keratinocytes and controls repair of skin wounds.^23,24^ However, all other type I IFNs are expressed upon stimulation by most nucleated cells.^25^ Early work suggested that the ability of IFN-α sub-types to enhance NK cell cytotoxicity varies over several orders of magnitude.^26^ More recently, several studies reported different potencies of type I IFNs in blocking viral infections and in exerting anti-tumour effects.^27–36^ Transcriptomic analysis indicates that different type I IFNs induce common and unique genes in CD4 T cells.^18^ These differences may be due to the observation that type I IFNs have varying characteristics of receptor binding.^37,38^ Combined with cell type-specific expression levels and stoichiometries of proteins involved in IFNAR signalling, different cell types have been suggested to respond in distinct ways to different type I IFNs. Indeed, mouse IFN-αA induces common and unique ISGs in different murine immune cells.^39^ Moreover, human B and CD4 T cells respond more strongly to IFN-α2 than to IFN-ω, while monocytes show equivalent responses.^37^ However, systematic studies comparing the effects of different type I IFNs in multiple types of primary cells are lacking at present.

Here, we close this knowledge gap using single cell technologies. We observed cell-type specific signalling and transcriptomic responses to type I IFN. Surprisingly, however, our comparison of multiple type I IFNs revealed only quantitative, but no apparent qualitative, differences. Our data mapping signalling and transcriptomic responses to type I IFNs represent a resource valuable to multiple scientific fields and are easily accessible through an interactive online interface (https://rehwinkellab.shinyapps.io/ifnresource/).

## Results

### Analysis of the signalling response to type I IFNs using mass cytometry

Systematic studies of the signalling pathways activated upon engagement of IFNAR with different type I IFNs in different cell types are currently lacking. To investigate this in multiple primary human cell types, we established a mass cytometry workflow to quantify cellular activation by phospho-protein staining in freshly isolated human peripheral blood mononuclear cells (PBMCs) stimulated with recombinant type I IFN (Figure 1b). We chose this model because of (i) the possibility of treating primary cells with minimal disturbance, (ii) the plethora of cell types found among PBMCs and ease of access to blood, (iii) the availability of bioactive human recombinant type I IFNs and (iv) the potential use of results as future biomarkers or for treatment strategies. We developed a 39 antibody staining panel comprising of 26 lineage markers to allow identification of approximately 20 major cell subsets and 13 antibodies to interrogate IFNAR signalling. The latter included antibodies against IFNAR1 and IFNAR2, nine phospho-proteins (pSTAT1, pSTAT3, pSTAT4, pSTAT5, pSTAT6, pERK1/2, pp38, pMAPKAPK2, pNFκB) and total STAT1 and STAT3. A suitable antibody for pSTAT2 was not available at the time this study began.

Phosphorylation of STAT proteins occurs rapidly and peaks around 30 minutes after IFNAR engagement.^12^ Here, we stimulated cells for 15 minutes to avoid saturation effects. To establish a suitable dose range, we stimulated PBMCs with 0, 25, 250, 2,500 or 25,000 U/ml of IFN-α2a and assessed the phosphorylation of STAT proteins. IFN-α2a was chosen because it is the IFN-α subtype most commonly used both *in vit*ro and *in vivo*. For dosing, we used bioactivity (U/ml), as determined by the manufacturer of the recombinant protein by inhibition of the cytopathic effect of encephalomyocarditis virus in A549 cells, instead of mass (pg/ml), because recombinant type I IFNs preparations may contain inactive protein. All STAT proteins were phosphorylated in response to IFN-α2a stimulation in a dose-dependent manner, saturating above the 2,500 U/ml dose (Figure 1c and d). STAT1 phosphorylation was the most sensitive response to stimulation, increasing even at the lowest dose used. In contrast, increased phosphorylation of the other STATs required stimulation with at least 250 U/ml IFN-α2a.

We next quantified the response to IFN-α2a in different cell types, taking advantage of our detailed phenotyping panel. Cells were clustered using the phenotyping markers and the resulting clusters were manually annotated, giving 17 distinct populations (Figure S1a and S1b). The expression of each pSTAT protein was then plotted for each cell population (Figure S1c, Supplementary Tables 1 and 2). The most abundant cell types (classical monocytes, naïve CD8+ T cells, CD56^lo^ NK cells, B cells and naïve CD4+ T cells) are shown in Figure 1e. Baseline levels of pSTAT1 varied between different cell types, with myeloid cells and central memory T cells having higher expression. Levels of pSTAT5 were highest in myeloid cells and plasmablasts at baseline whereas pSTAT3, 4 and 6 were undetectable in unstimulated cells (Figure S1 and S2). STAT1 was phosphorylated in all cell types, in line with its role in canonical signalling downstream of IFNAR. STAT3, STAT5 and STAT6 phosphorylation was also detected in all cell types, although only at concentrations of 250 U/ml IFN-α2a and above. In contrast, STAT4 phosphorylation only occurred in some T cell subsets and NK cells, reflecting the largely T cell and NK cell restricted expression of STAT4.^40^ This provides an explanation for the lower pSTAT4 response we observed when we analysed all cells together (Figure 1c). Notably, there was considerable heterogeneity in the extent of STAT phosphorylation within each cell population (Figure 1e and S1c).

These data show that our mass cytometry workflow successfully and simultaneously quantified phosphorylation of multiple proteins in response to stimulation with type I IFN. Our results further highlight that different types of cells respond in different ways to type I IFN, and that cell type specific responses may be masked by bulk analysis.

### Different type I IFNs induce qualitatively similar signalling responses in PBMCs

We next used our mass cytometry workflow to investigate the response to different type I IFNs. Cells were stimulated for 15 minutes with 2,500 U/ml of commercially available bioactive recombinant protein for the 13 IFN-α subtypes, IFN-β and IFN-ω. IFN-ε and IFN-κ were not included as their expression is restricted to specific tissues (female reproductive tract and skin, respectively).^21,23^ The concentration was chosen to ensure that phosphorylation events of all STAT proteins could be detected. All type I IFNs induced phosphorylation of the STAT proteins when all cells were analysed together (Figure 2a and 2b). Analysis of individual cell populations revealed that, for any given cell type, all type I IFN subtypes led to phosphorylation of STAT1 to a similar extent, although clear differences in pSTAT1 levels were evident between different cell types (Figure 2c and 2d, Supplementary Tables 3 and 4). For pSTAT3 (Figure 2e), pSTAT4 (Figure 2f), pSTAT5 (Figure 2g) and pSTAT6 (Figure 2h), quantitative differences were evident between type I IFN subtypes. For example, IFN-α1 induced lower levels of pSTAT4 and IFN-α1, -α2b and -α14 induced lower levels of pSTAT6. However, qualitative differences in the response to different type I IFN subtypes were not apparent amongst the cell types analysed. Similar results were obtained for phosphorylation of NFκBp65, p38, MAPKAPK2 and ERK1/2 (Figure S3).

**Figure 2.**
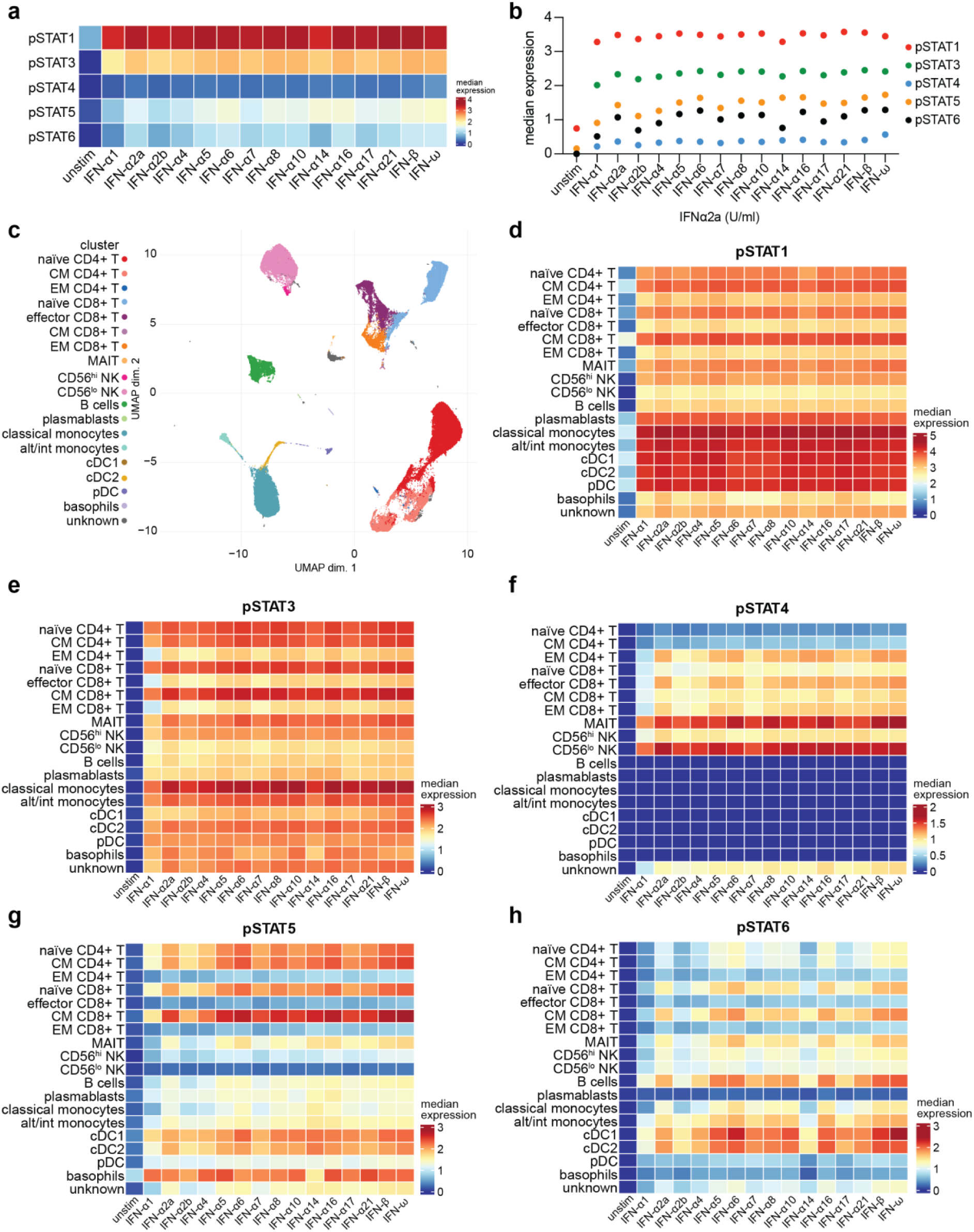
Phosphorylation of STAT proteins in response to different type I IFNs. (a) Median expression of pSTATs in PBMCs in response to treatment with 2500 U/ml of the indicated type I IFNs for 15 minutes. (b) Depiction of the data shown in (a) as a dot plot. (c) UMAP plot showing clustering of PBMCs and identification of different cell types based on expression of the phenotyping markers. (d – h) Heatmaps showing median expression of pSTATs in each cell type in response to treatment with different type I IFNs. Data are from one experiment with one donor. See also Figures S3 – S6 and Supplementary Tables 3 – 8.

To investigate the effects of different type I IFNs on phosphorylation of STAT proteins at later time points, we first stimulated PBMCs for 90 minutes with increasing doses of a subset of five type I IFNs (IFN-α1, -α2a and -α10; IFN-β; IFN-ω). These represent low, medium and high potency IFN-α subtypes (Figure 2), together with the unique IFN-β and IFN-ω. At this time point, phosphorylation levels of all STATs were lower compared to the 15 minute timepoint, with only small increases even at 2,500 U/ml (Figure S6). Quantitative differences in the amount of each type I IFN required to increase STAT phosphorylation were evident, with higher doses of the three IFN-α subtypes required compared to IFN-β and IFN-ω (Figure S4). Increased baseline levels of pSTAT1 and pSTAT3 were seen at this timepoint (Figure S6), likely a consequence of the increased time cells spent in culture before analysis. Stimulation with type I IFNs increased phosphorylation of STAT1 in all cell types; however, as observed 15 minutes after stimulation, differences between type I IFN subtypes were largely quantitative (Figure S4d, Supplementary Tables 5 and 6). Only slight increases in phosphorylation of the other STAT proteins were evident at this time point and again differences between type I IFN subtypes were quantitative (Figure S4e-h). Similar results were obtained when PBMCs were stimulated for 24 hours (Figure S5, Supplementary Tables 7 and 8).

Taken together, these data show that in PBMCs stimulated *ex vivo*, different recombinant type I IFNs induce similar profiles of transcription factor phosphorylation, although the magnitude of the response varies between type I IFNs.

### Different type I IFNs induce qualitatively similar transcriptional responses in PBMCs

IFNAR signalling profoundly alters the transcriptome. In addition to the induction of ISGs, type I IFNs may downregulate mRNA levels of some genes.^17,18^ We stimulated PBMCs from three donors with 250 U/ml type I IFN, using the set of five type I IFNs described above. 24 hours after stimulation, we extracted RNA and, following enrichment of polyadenylated transcripts, performed RNA sequencing. The 24 hour timepoint was chosen based on RT-qPCR analysis of the induction of seven ISGs over time in response to IFN-α1 and IFN-β. Although the levels of some ISGs peaked 2-4 hours after stimulation, these transcripts were still upregulated after 24 hours, a timepoint at which other ISGs were maximally induced (Figure S7). Therefore, the 24-hour timepoint likely allows detection of early and late responding ISGs. Sequencing reads were aligned to the human transcriptome (Figure S8a, S8b, Supplementary Table 9). We confirmed that cells from all three donors had comparable expression levels of genes encoding the key components of IFNAR signalling (Figure S8c). Next, we performed differential gene expression analysis for cells stimulated with each type I IFN compared to the unstimulated samples. Many protein-coding genes were differentially expressed in response to each type I IFN, with more differentially expressed genes (DEGs) up-regulated than down-regulated (Figure 3a, Supplementary Tables 10 and 11). IFN-β induced the largest number of DEGs and IFN-α1 the least. On the whole, genes that were upregulated showed the greatest fold change and significance (Figure 3a, Figure S8d, Figure S9). Using a low stringency filter for fold change (1.5), we identified 2,430 genes that were up-regulated and 2,130 that were down-regulated in response to at least one type I IFN (Figure 3c). Of these, 638 were significantly up-regulated by all type I IFNs and 82 significantly down-regulated (Figure S10a).

**Figure 3.**
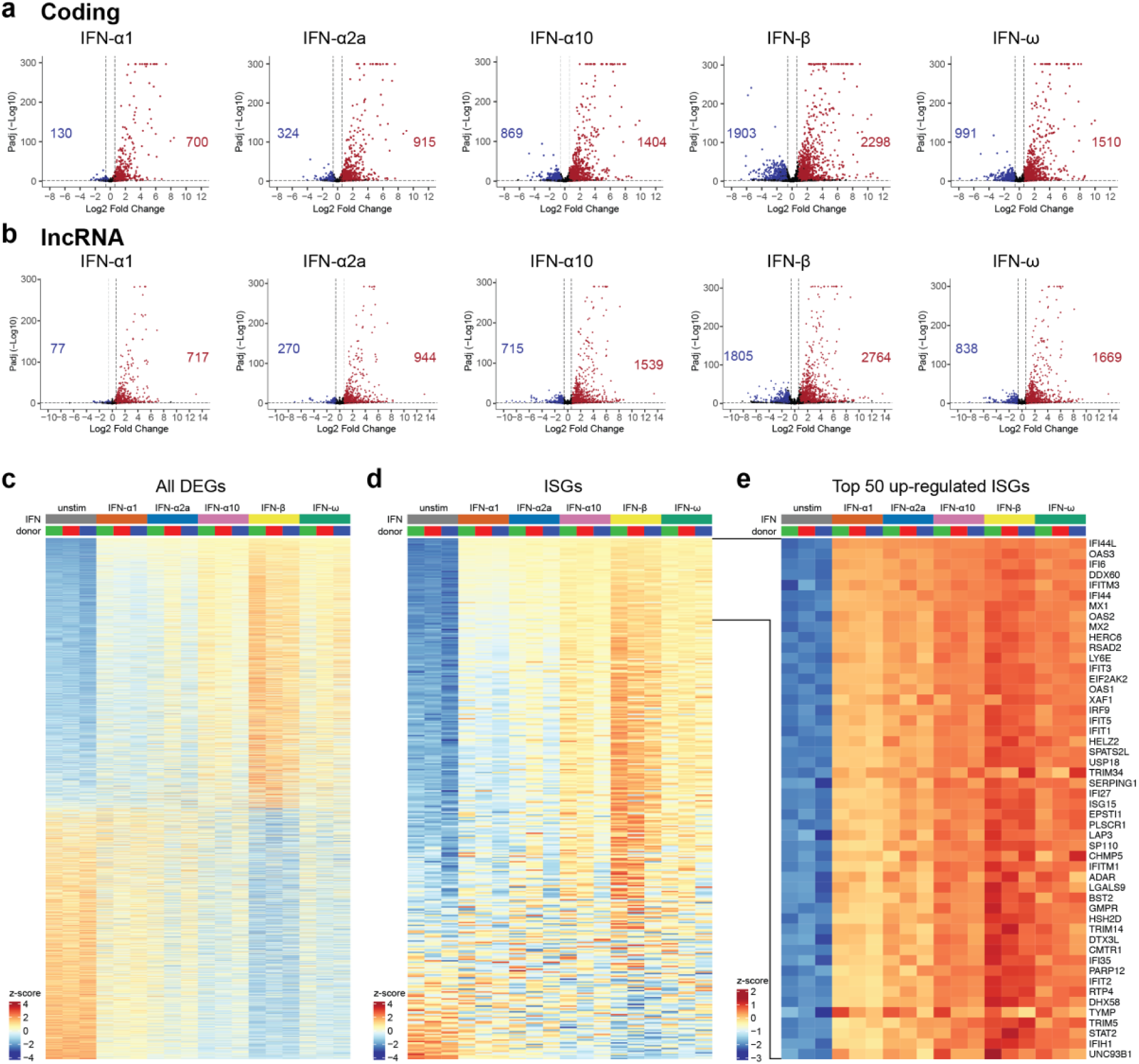
Bulk RNAseq of type I IFN-stimulated PBMCs. (a,b) Volcano plots of differentially expressed protein-coding genes (a) and lncRNAs (b) in PBMCs stimulated with 250 U/ml of the indicated type I IFNs for 24 hours, compared to mock stimulated PBMCs. Red indicates significant upregulation of expression (padj < 0.05, fold change > 1.5) and blue indicates significant downregulation of expression (padj < 0.05, fold change < -1.5); the corresponding numbers of genes are shown. (c) Heatmap of all protein-coding genes that are significantly differentially expressed in response to at least one type I IFN, compared to unstimulated PBMCs (2,430 up-regulated and 2,130 down-regulated genes). (d, e). Heatmaps of expression of 379 annotated ISGs (d) and top 50 up-regulated ISGs (e). In c - e, genes are ranked by z-score of all type I IFN-stimulated samples. Data are from one experiment with three donors. See also Figures S7 – S12 and Supplementary Tables 9 – 15.

Gene Ontology (GO) analysis for biological processes associated with DEGs regulated by all type I IFNs confirmed that, as predicted, GO terms associated with the response to type I IFNs and the antiviral response were most enriched among the up-regulated DEGs (Figure S11a). Using a published list of 379 human ISGs compiled from microarray data from multiple cell lines and tissues^16^ (Supplementary Table 12), we confirmed that 250 were up-regulated by at least one IFN in PBMCs (Figure 3d), with classical ISGs such as *IFI44L*, *MX1* and *IFIT3* among the most up-regulated (Figure 3e). PBMCs from all three donors showed comparable induction of most ISGs. Many of the most significantly induced genes were known ISGs (Figure S8d), expression of which was often undetectable in unstimulated PBMCs.

In addition to inducing the expression of many genes, type I IFNs also significantly repressed the expression of another set of genes (Figure 3c). We used GO analysis to gain insight into biological processes related to the 82 genes significantly downregulated by all type I IFNs (Figure S11b). This analysis did not reveal enrichment of GO categories containing more than a few type I IFN-repressed genes.

We also considered genes that met our criteria for significance (padj < 0.05, 1.5 fold up- or down-regulated) for one type I IFN only. The vast majority of these were found in response to IFN-β (Figure S9). Visualisation of the expression of these genes across all samples using heatmaps suggested that, for the vast majority of these genes, the differences in expression between the different type I IFNs was quantitative (Figure S10). For example, genes that were only upregulated above threshold for IFN-β stimulation showed a clear trend towards induction by the other four type I IFNs tested, albeit to lower levels (Figure S10).

### Expression of lncRNAs in PBMCs following stimulation with type I IFNs

In addition to protein coding genes, we also investigated the effect of type I IFN stimulation on the expression of long noncoding RNAs (lncRNAs). The majority of lncRNAs are polyadenylated^41^; therefore, we were able to analyse their expression in our bulk RNAseq dataset. We aligned the sequencing reads to a reference database of 127,802 transcripts (LNCipedia) using kallisto.^42^ An average of 12 million reads/sample aligned to this database (Figure S8b), with 57,812 transcripts passing the filter of having > 3 counts and being expressed in 2 or more samples. Differential gene expression analysis for each of the type I IFN stimulated samples compared to the unstimulated sample showed that, similar to the expression of protein-coding genes, type I IFNs both induced and repressed expression of lncRNAs. More lncRNAs were up-regulated than down-regulated (Figure 3b, Figure S8e, Figure S11, Supplementary Tables 13 - 15). 628 lncRNAs were significantly up- and 46 down-regulated by all type I IFNs (padj < 0.05, fold change of 1.5) (Figure S12b).

Many lncRNAs up-regulated in response to treatment with type I IFNs are located close to protein-coding ISGs, and may act to regulate their expression.^43^ In the absence of their own official gene symbol, lncRNAs are named after the gene of the nearest protein-coding gene on the same strand. We therefore used the list of protein-coding ISGs to investigate the extent to which lncRNAs located near ISGs were differentially regulated by type I IFN in our dataset. Of the 674 lncRNAs that were significantly differentially expressed in response to all five type I IFNs tested (padj < 0.05, fold change of at least 1.5, Figure S12c), 58 were ‘associated’ with ISGs (56 up-regulated, 2 down-regulated) (Figure S12d).

Together, these data show that while a robust ISG response was induced in PBMCs following 24 hours of stimulation with type I IFN, there were only minimal differences between type I IFN subtypes.

### Analysis of the ISG response to type I IFNs in PBMCs at the single cell level

PBMCs represent a heterogeneous mix of multiple different cell types of varying frequencies. Given that different cell types displayed different signalling responses to type I IFN stimulation (Figures 1 and 2) and the possibility of cell-to-cell variation, we used scRNAseq to further analyse transcriptional responses to type I IFNs. We treated PBMCs from one donor with 250 U/ml IFN-α1, -α2a, -α10, -β or -ω for 24 hours. We took advantage of the utility of our mass cytometry phenotyping panel by staining cells with CITE-seq antibodies corresponding to 20 of these markers prior to capture on a 10X Genomics platform. In brief, CITE-seq involves antibodies conjugated to nucleic acid-based barcodes with a polyA tail, which are sequenced alongside mRNA from captured single cells.^44^ After demultiplexing and quality control, the dataset consisted of 19,587 single cells, with an approximately equal split between samples (unstimulated = 2,690 cells, IFN-α1 = 3,328, IFN-α2a = 3,426, IFN-α10 = 3,332, IFN-β = 3,350, IFN-ω = 3,461). Seurat Weighted Nearest Neighbor (WNN) analysis was used to cluster cells based on mRNA and protein expression^45^ and cells were visualised using UMAP (wnnUMAP) (Figure 4a, Figure S13). Cell types were identified based on expression of protein markers, in combination with mRNA expression of known markers not present in the CITE-seq antibody panel. A number of markers were detected equally well at protein and mRNA levels, such as CD3, but mRNAs for other markers including CD19, CD11c and CD4 were not detectable (Figure 4b).

**Figure 4.**
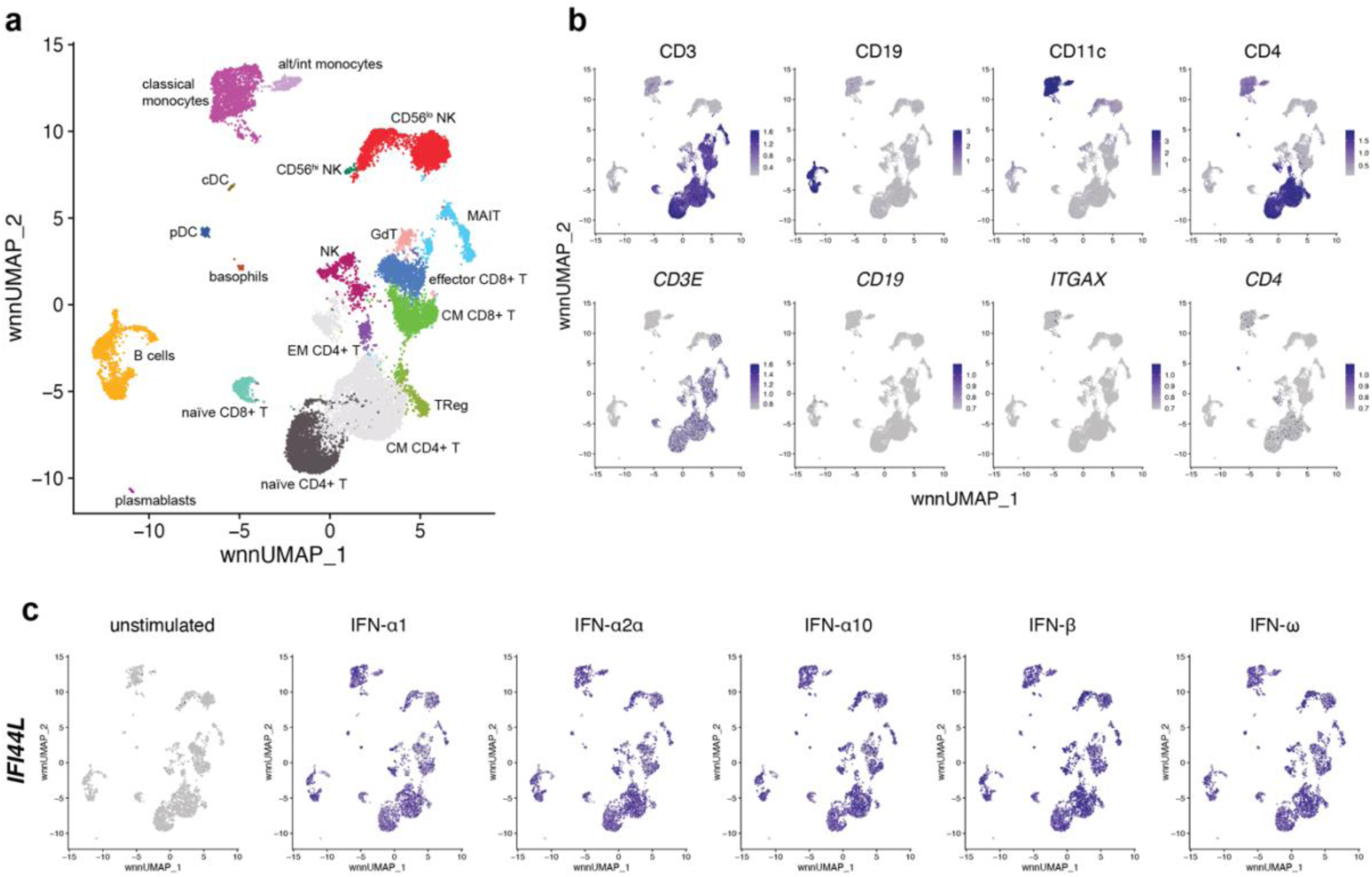
Single cell RNAseq analysis of PBMCs stimulated with type I IFNs. (a) UMAP visualisation of 19,587 single cells clustered following weighted nearest neighbour (WNN) analysis integrating protein and RNA data for each cell. (b) UMAP plots showing expression of the phenotyping markers CD3, CD19, CD11c and CD4 at the protein (top row) and RNA (bottom row) level. (c) UMAP showing expression of *IFI44L* in unstimulated and type I IFN-stimulated samples. Data are from one experiment with one donor. See also Figures S13 and S14 and Supplementary Tables 16 – 22.

To identify genes induced in response to stimulation with each type I IFN, we performed differential expression analysis for each cell type, comparing type I IFN-treated samples to the unstimulated sample (Supplementary Tables 18 - 22). This was only possible for cell types with an average of > 50 cells per sample, precluding this analysis for pDCs, basophils, CD56^hi^ NK cells, cDCs and plasmablasts. We identified only 10 genes that were significantly up-regulated in all cell types in response to all five type I IFN subtypes tested here: *IFI44L*, *ISG15*, *IFIT3*, *XAF1*, *MX1*, *IFI6*, *IFIT1*, *TRIM22*, *MX2* and *RSAD2* (Figure 4c, Figure S14). We term these genes that universally mark the type I IFN response ‘Core ISGs’.

### Cell type-specific ISG

Next, we investigated whether ISGs were regulated in a cell type-specific manner. We first established how many genes were significantly up-regulated by monocytes (classical monocytes, alternative/intermediate monocytes) and lymphocytes (central memory CD4+ T cells, naïve CD4+ T cells, CD56^lo^ NK cells, B cells, central memory CD8+ T cells, effector CD8+ T cells, MAIT cells, other NK cells, naïve CD8+ T cells, regulatory T cells, effector memory CD4+ T cells and γδ-T cells) in response to all type I IFNs. 225 genes were up-regulated by monocytes and 175 by lymphocytes, with 107 genes significantly up-regulated by both using padj < 0.05 and 1.5 fold induction as thresholds (Figure 5a, Figure S15). Genes specifically up-regulated by monocytes included *IFITM3*, *CXCL10* and *SIGLEC1* while genes specifically up-regulated by lymphocytes included *LAG3*, *ALOX5AP* and *CD48* (Figure 5b and c). As expected, genes up-regulated by both monocytes and lymphocytes included those described as ISGs by many previous studies, such as *ISG15*, *IFI44* and *IFIT5* (Figure 5b and c). Although these genes were selected based on their up-regulation by all five type I IFN subtypes tested, differences in the magnitude of the response were apparent, with IFN-β inducing many ISGs most strongly. This was in agreement with the mass cytometry and bulk RNAseq data indicating that different type I IFNs induce qualitatively similar responses. Subdivision of lymphocytes into T cells, B cells and NK cells revealed 40 genes and two lncRNAs only up-regulated in T cells, six genes and one lncRNA only up-regulated by B cells and two genes that were only up-regulated by NK cells (Figure S16) using our significance and fold-change thresholds.

**Figure 5.**
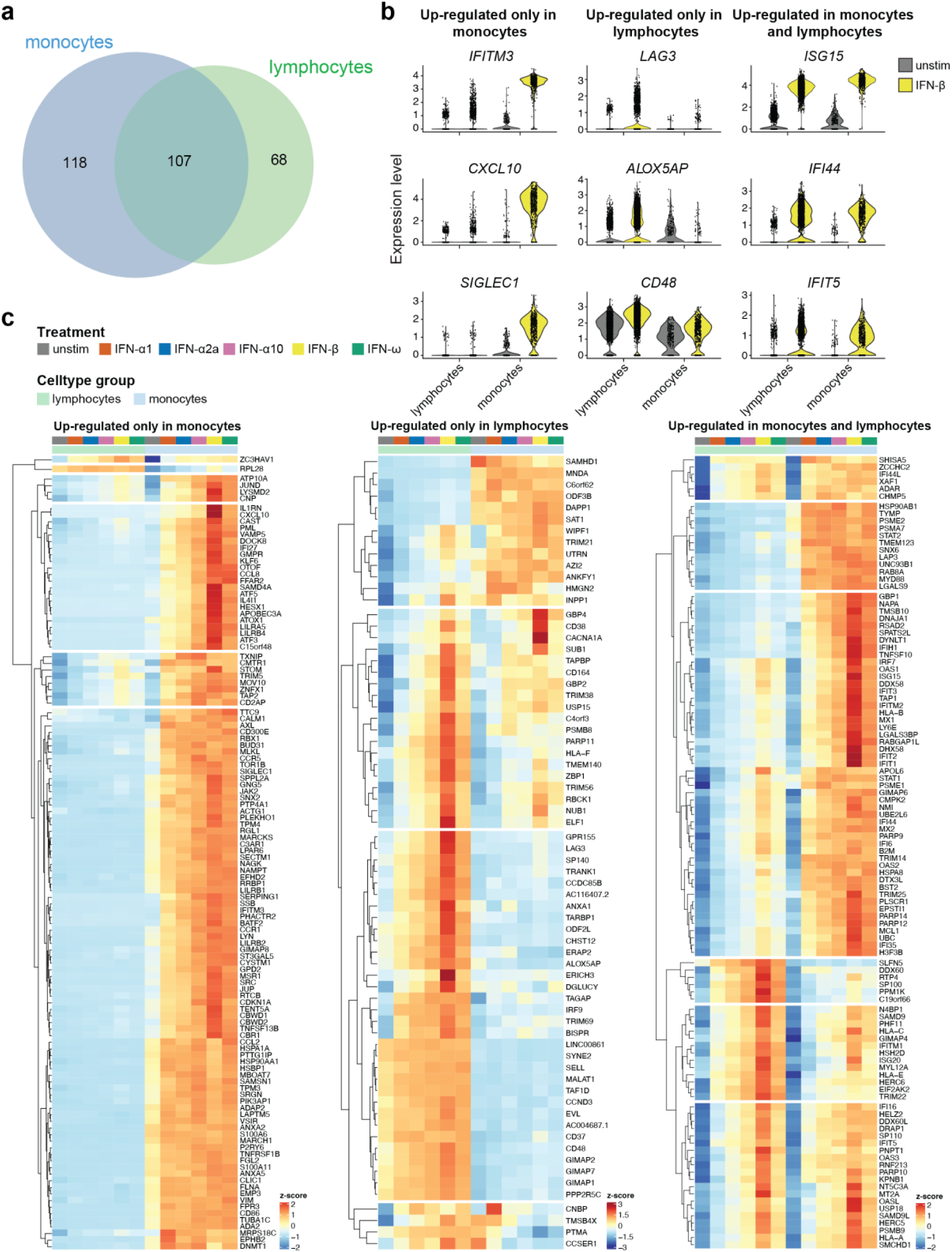
Cell type-specific responses to type I IFNs. (a) Venn diagram showing the number of genes significantly up-regulated in response to all tested type I IFNs in monocytes and lymphocytes. Gene lists are provided in Figure S15. (b) Violin plots showing expression of selected genes in unstimulated or IFN-β-treated cells. (c) Heatmaps showing average expression of significantly up-regulated genes for each sample. Expression is represented as a z-score across all samples for each gene. Data are from one experiment with one donor. See also Figures S15 – S17.

In the bulk RNAseq dataset, differences in the expression of genes induced by the different type I IFN subtypes were largely quantitative (Figure S10). To investigate if this was also the case when cell type-specific ISGs were taken into account, we analysed the expression of genes induced only by one type I IFN subtype in monocytes and lymphocytes in all samples (Figure S17). Consistent with our bulk RNAseq data, differences were again quantitative rather than qualitative for most genes.

### Heterogeneity of the response to type I IFNs

Next, we asked whether there was, for a given cell type, cell-to-cell variation in the transcriptional response to type I IFN stimulation. For example, some cells may be “super-responders” and others “non-responders”. It is also possible that some cells up-regulate a distinct module of ISGs in response to type I IFN while others up-regulate a different set of ISGs, information that would be masked when averaging across the cell type cluster.

We chose to focus on classical monocytes as these represent a large cluster with many significantly up-regulated genes (i.e., 278, 330, 482, 643, 526 and 225 in response to IFN-α1, -α2a, -α10, -β, -ω and all five type I IFNs, respectively). Expression of each gene for each cell within this cell type was visualised using heatmaps and the genes clustered using Euclidean distance. This is shown for unstimulated and IFN-β-treated cells in Figure 6a (see Figure S18 for this heatmap with gene name annotations for online viewing). This revealed that there was a large spectrum of ISG induction in IFN-β-stimulated cells, with some genes induced moderately (such as those in cluster 2, including classical ISGs such as *IFI44*, *OAS2*, *OAS3* and *OASL*), intermediately (clusters 4 and 6) and strongly (cluster 5). The genes in cluster 5 were all expressed at very low level in the unstimulated cells and therefore showed the greatest fold induction in expression upon stimulation with IFN. There were also differences in baseline expression of many of these ISGs, with most being not expressed/expressed at very low levels by most cells. Most of the 225 genes induced by all type I IFNs tested were up-regulated by all cells in the IFN-β-treated sample compared to the unstimulated cells. Interestingly, some genes displayed heterogeneous expression in the IFN-β-treated sample (Figures 6a and S18) and in the samples stimulated with other type I IFNs (Figure S19). These included *IFI27*, *CCL8, CCL2* and *CXCL10* (Figure S18). To establish if expression of these four genes was linked, i.e. whether cells which up-regulated one of them also up-regulated the others, we clustered IFN-β stimulated classical monocytes based on their expression of these genes (Figure S20). This revealed that whilst there were cells that up-regulated all four genes, there were also cells which expressed three or fewer, in all combinations, with no obvious patterns. This suggests it is unlikely that these four ISGs were induced in relation to each other.

**Figure 6.**
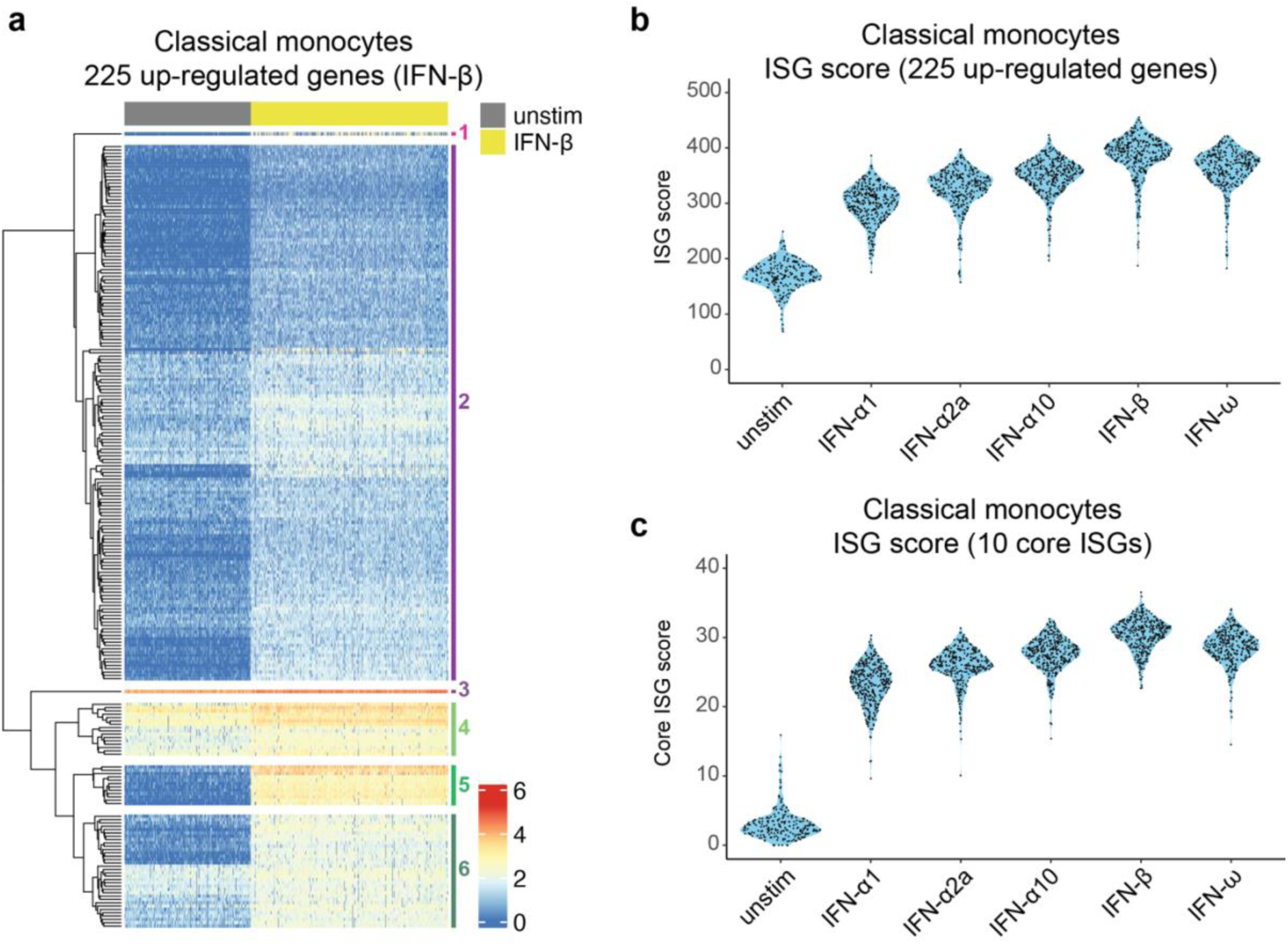
Response of classical monocytes to type I IFNs. (a) Heatmap showing expression of the 225 genes significantly up-regulated in response to all tested type I IFNs in classical monocytes in the unstimulated and IFN-β-stimulated conditions. Each row represents a gene and each column a cell. An enlarged heatmap with annotated gene names is provided in Figure S18. (b,c) Sinaplots with violin outline showing the ISG scores calculated using the 225 up-regulated genes in classical monocytes (b) and our list of ten core ISGs (c). ISG scores were determined on a per cell basis and are shown for classical monocytes with each black dot representing a single cell. Data are from one experiment with one donor. See also Figures S18 – S26.

We also visualised expression of this list of 225 genes up-regulated by classical monocytes in B cells and naïve CD4+ T cells, two clusters with comparable numbers of cells (Figure S21). As expected, a large number of these genes were unchanged in these cell types in response to type I IFN-treatment. Similarly, with the list of 94 genes up-regulated by B cells (Figure S22) and 113 genes up-regulated by naïve CD4+ T cells (Figure S23), cell type-specific responses were clear. Genes that were up-regulated in all three cell types showed a similar degree of heterogeneity between individual cells (Figure S24).

Finally, we calculated an ISG score for every cell in our dataset by summing the log transformed expression values for each gene significantly up-regulated by the corresponding cell type. As shown for classical monocytes in Figure 6b, the distribution of the scores for cells within a sample indicated that there are no “super-responder” cells and very few “non-responders”. This was also the case for other cell types (Figure S25). We also calculated a Core ISG score for each cell using the ten ‘Core ISGs’ (Figure 4c and S14). This confirmed that all cells within a stimulated sample up-regulated expression of these genes, regardless of the type I IFN subtype used or cell type (Figure 6c and Figure S26). We therefore propose that an ISG score calculated from expression of these ten genes is sufficient to capture a transcriptional response to all type I IFN subtypes in all types of PBMCs.

Taken together, our single cell transcriptomic analysis of type I IFN stimulated PBMCs identified ISGs induced in different white blood cells. Some ISGs were induced broadly in all or many cell types, while others displayed cell-type specific induction. In agreement with the mass cytometry and bulk RNAseq data, our scRNAseq analysis also suggested that different type I IFNs induced qualitatively similar responses.

## Discussion

Type I IFNs are a large family of cytokines, playing important roles in the immune system and in many diseases that range from infections to inflammatory and malignant conditions. Here, we employed single-cell technologies to generate comprehensive maps of signalling and transcriptomic changes in human white blood cells stimulated *ex vivo* with different type I IFNs. These data revealed responses shared between cell types as well as cell-type specific behaviours. Moreover, when comparing different type I IFN subtypes, we observed qualitatively similar responses that varied in magnitude.

Our datasets are freely available in multiple ways. Raw data can be accessed through repositories (see data availability statement). Lists of differentially expressed genes in different cell types in response to different type I IFNs are provided in the supplement to this article. In addition, we provide our data via the Interactive SummarizedExperiment Explorer (iSEE).^46^ This online tool [https://rehwinkellab.shinyapps.io/ifnresource/] is easy to use and requires no programming skills. Examples of its application include (1) querying a cell type of interest for genes induced or repressed by type I IFNs or for its signalling response and (2) analysing a gene (or signalling protein) of interest for baseline and type I IFN-regulated expression (or phosphorylation) in multiple populations of leukocytes. Another widely applicable result is the list of ten ‘Core ISGs’ induced in all types of PBMCs by all type I IFNs. For example, this set of ISGs will be useful as a biomarker to test for the presence of a type I IFN response in patients where the cellular basis of disease in unknown.

Monocytes were the cell type with the largest number of type I IFN-regulated genes, both in terms of the overall set of differentially expressed genes as well as genes affected only in monocytes but not in other cell types. This was likely due in part to the comparatively high baseline expression and/or phosphorylation of proteins involved in IFNAR signalling in monocytes (Figure S2), bestowing an enhanced capacity to respond. Another important aspect likely determining which genes are regulated by type I IFN in different cell types is the epigenetic landscape. In the future, it would be interesting to investigate whether the contingent of ISGs in each cell type correlates with chromatin marks and accessibility using approaches such as ChIPseq and ATACseq.

In addition to protein-coding genes, we also provide a systematic analysis of lncRNAs using our bulk RNAseq data. lncRNAs are a heterogenous group broadly defined as transcripts > 200 nt in length that do not encode proteins.^47^ They are generally less evolutionarily conserved than protein-coding genes, often with tissue-specific expression, and exhibit a wide range of cellular functions.^48^ The expression of many lncRNAs is regulated and alterations to their expression can be associated with human diseases. Several lncRNAs are associated with the antiviral response and regulate the activity of pattern recognition receptors and signalling pathways leading to cytokine production, including the type I IFN signalling pathway.^43,49^ These included NRIR (Negative Regulator of the Interferon Response)^50^, previously implicated in driving a type I IFN signature in monocytes in human autoimmune disease and viral infection.^51–53^ Indeed, NRIR was differentially expressed in our dataset (Figure S12g). Here, we defined large numbers of lncRNAs induced and repressed in PBMCs, some of which were encoded near ISGs. These data provide a rich resource for future studies investigating functional roles of interferon-stimulated or -repressed lncRNAs.

An important and open question in the type I IFN field pertains to the large number of genes encoding these cytokines. Many mammalian species, including bats, encode a multitude of type I IFNs.^54–56^ This is indicative of evolutionary pressures leading to maintenance of many type I IFN genes. Simply put, why are there so many type I IFNs all using the same receptor?

One possible explanation is that different type I IFNs have unique activities against specific pathogens or in specific cell types.^26–36^ For example, IFN-α14 is particularly potent at inhibiting HIV-1 replication in different *in vitro*, *ex vivo* and *in vivo* models compared to IFN-α2.^27–29,34^ During SARS-CoV-2 infection, IFN-α5 exerts a strong antiviral effect in cultured human airway epithelial cells.^33^ The latter observation correlates with induction of a subset of ISGs by IFN-α5 and other potently antiviral type I IFNs; these ISGs are not induced in this setting by other type I IFNs with poor anti-SARS-CoV-2 activity.^33^ In contrast, we observed in PBMCs that different type I IFNs did not trigger such qualitatively different cellular behaviours. Instead, only the magnitude of the signalling and transcriptional responses varied between different type I IFNs. These divergent observations may be due to differences in haematopoietic and non-haematopoietic cells. For example, it is possible that in airway epithelial cells the stoichiometries and activities of proteins involved in IFNAR signalling are different to white blood cells and that this, perhaps in conjunction with alternate chromatin configurations, permits type I IFN-subtype-specific responses due to differences in their affinities for IFNAR. Studies comparing cells representing different tissues are therefore warranted, as is careful dissection of the responses to type I IFNs over varying doses and at different time points. It is tempting to speculate that additional factors such as type I IFN ‘presentation’ involving neighbouring cells in solid tissues or the soluble IFNAR2 isoform^57^ in blood may play currently under-appreciated roles in controlling responses to type I IFNs.

Other possible explanations for the multitude of type I IFNs include pathogen encoded antagonists. Indeed, vaccinia virus encodes B18, a secreted protein that sequesters type I IFNs^58^. Expansion of the type I IFN locus may have evolved to overcome the effects of B18 and potentially of similar, yet to be discovered, inhibitors, either simply by increasing gene dosage to saturate antagonists or more specifically through sequence diversification allowing escape of inhibition. Additionally, or alternatively, differences in the kinetics and signalling requirements for induction of type I IFNs may explain the large number of these cytokines. For example, the IFN-β promoter has elements recognised by multiple transcription factors including IRF3 and IRF7, NFκB and AP1 family members. In contrast, IFN-α genes are only controlled by IRF3 and IRF7^59^. Moreover, the different IFN-α genes are induced at different time points during viral infection^60^. It is therefore possible that so many type I IFN genes have evolved to allow temporal, and perhaps also spatial and cell-type specific, control of their induction, including feedback loops. Indeed, IRF7 is encoded by an ISGs and is an important part of a positive feed-forward circuit promoting strong IFN-α production by plasmacytoid dendritic cells^61,62^. This scenario would be consistent with our findings that type I IFNs – once produced – are functionally equivalent. Clearly, understanding the diversity of type I IFNs remains an important challenge for future studies.

Taken together, we provide a data-rich and easily accessible resource of the responses of many different types of primary immune cells to type I IFNs. We anticipate that our datasets will instruct and inform research in many fields ranging from immunology and virology to cancer.

### Limitations of the study

This study was performed in PBMCs stimulated with type I IFNs *ex vivo*. Other cell types may show different behaviours in response to type I IFNs and additional factors such as the soluble form of IFNAR2 may impact *in vivo* responses. In-depth analysis of some cell types, such as rare dendritic cell subsets, was not possible due to the small number of cells. Enrichment of these cells prior to analysis will be necessary in future studies. We used recombinant type I IFNs for treatment. Doses were normalised using bioactivity (U/ml), determined by inhibition of the cytopathic effect of encephalomyocarditis virus in A549 cells. It is possible that the antiviral effect in this setting depends on induction of one or few ISGs, rather than on the breadth of effects of type I IFNs, potentially skewing normalisation. However, other ways of normalisation, for example using mass (pg/ml), have inherent limitations, too, because recombinant type I IFN preparations likely contain inactive protein.

## Methods

### Study subjects

PBMCs were isolated from the peripheral blood of healthy donors using Lymphoprep (Stemcell Technologies), according to the manufacturer’s instructions.

### Data acquisition

#### Mass cytometry (CyTOF)

PBMCs were washed in serum-free RPMI and then resuspended at 10^7^ cells/ml in serum-free RPMI containing 0.5 mM Cell-ID Cisplatin (Fluidigm) and incubated at 37°C for 5 min. Cells were washed with RPMI containing 10% (v/v) FCS (Sigma) and 2 mM L-Glutamine (R10), centrifuged at 300 x *g* for 5 min before being resuspended to 6 x 10^7^ cells/ml in R10 and rested at 37°C for 15 min. 50 μl of cells (3 x 10^6^) cells were transferred to 15 ml falcon tubes for stimulation and antibody staining. Antibody staining was performed as described previously^63^. Antibodies are listed in the Key Resources Table. Staining for CD14, CCR6, CD56, CD45RO, CD27, CCR7, CCR4 and CXCR3 was done for 30 min in R10 at 37°C, prior to stimulation/fixation as epitopes recognised by these antibodies are sensitive to fixation. Cells were stimulated with the indicated concentration of type I IFN (Supplementary Table 2) diluted in R10 for the indicated time at 37°C. After washing with 5 ml Maxpar PBS (Fluidigm), cells were fixed with 1 X Maxpar Fix I Buffer (Fluidigm) for 10 min at room temperature before being washed with 1.5 ml Maxpar Cell Staining Buffer (CSB, Fluidigm). All centrifugation steps after this point were at 800 x *g* for 5 min. Cells were barcoded using Cell-ID 20-Plex Pd Barcoding Kit (Fluidigm), according to the manufacturer’s instructions, and washed twice with CSB before samples were pooled and counted. All further steps were performed on pooled cells. Fc receptors were blocked using Fc Receptor Binding Inhibitor Antibody (eBioscience), diluted 1:10 in CSB for 10 min at room temperature. Surface antibody staining mixture was added directly to the blocking solution and incubated for 30 min at room temperature. Cells were washed twice with CSB, resuspended in ice-cold methanol and stored at -80°C overnight. After washing twice with CSB, cells were stained with intracellular antibody staining mixture for 30 min at room temperature before two further washes in CSB. Cells were resuspended in 1.6% (v/v) formaldehyde (Pierce, #28906) diluted in Maxpar PBS and incubated for 10 min at room temperature. Cells were resuspended in 125 mM cell-ID Intercalator (Fluidigm) diluted in Maxpar Fix and Perm Buffer (Fluidigm) and incubated overnight at 4°C. Compensation beads (OneComp eBeads Compensation Beads, Invitrogen, #01-111-42) stained with 1 μl of each antibody were also prepared. The following day, cells and compensation beads were washed twice with CSB and twice with Maxpar water (Fluidigm), mixed with a 1:10 volume of EQ Four Element Calibration Beads (Fluidigm) before acquisition on a Helios Mass Cytometer (Fludigm) using the HT injector.

#### qRT-PCR

Freshly isolated PBMCs were stimulated with 250 U/ml of IFN-α1 or IFN-β in R10 or left unstimulated (3 x 10^6^ cells/sample) in a volume of 2 ml in a non-tissue culture treated 12 well plate and incubated for the time indicated in the figure legend at 37°C. RNA was extracted using a RNeasy Mini Plus Kit including a gDNA eliminator column step, according to the manufacturer’s instructions. cDNA synthesis was performed with SuperScript IV reverse transcriptase (Thermo Fisher Scientific) and oligo(dT)_12-18_ primers (Thermo Fisher Scientific). 15 ng of cDNA was amplified using Taqman Universal PCR Mix (Thermo Fisher Scientific) and Taqman probes (see Key Resources Table) in 5 μl reactions. qPCR was performed on a QuantStudio 7 Flex real-time PCR system (Applied Biosystems). mRNA expression data were normalised to *HPRT* and analysed by the comparative C_T_ method. Fold changes in expression from unstimulated samples for each time point were calculated for the IFN-treated samples.

#### Bulk RNA-seq

Freshly isolated PBMCs were stimulated with 250 U/ml of each type I IFN in R10 (3 x 10^6^ cells/sample) in a volume of 2 ml in a non-tissue culture treated 12 well plate and incubated for 24 h at 37°C. Cells were harvested by gentle pipetting and centrifuged at 300 x g for 5 min. The pellet was resuspended in 350 μl RLT buffer from an RNeasy Plus Mini Kit (QIAGEN) and transferred to a QiaShredder column. After homogenisation by centrifuging at 16,000 x g for 2 min, RNA was extracted according to the manufacturer’s instructions, including a gDNA eliminator column step. RNA was quantified using a Qubit 3.0 RNA BR (broad-range) Assay Kit (Invitrogen) and using RiboGren (Invitrogen) on a FLUOstar OPTIMA plate reader (BMG Labtech). The RNA size profile and integrity was assessed using a Tapestation 4200 with High Sensitivity RNA ScreenTape (Agilent Technologies). Input material was normalised to 100 ng prior to library preparation by the Oxford Genomics Centre. Polyadenylated transcript enrichment and strand specific library preparation was completed using a NEBNext Ultra II mRNA kit (New England Biolabs) following the manufacturer’s instructions. Libraries were amplified (17 cycles) using in-house unique dual indexing primers^64^. Individual libraries were normalised using a Qubit, and the size profile was analysed using a Tapestation before individual libraries were pooled. The pooled library was diluted to ∼10 nM, denatured and further diluted prior paired-end sequencing (150 bp reads) on a NovaSeq 6000 platform (Illumina, NovaSeq 6000 S2/S4 reagent kit, 300 cycles), yielding between 38 – 55 million reads/sample.

#### scRNAseq: Cell stimulation and capture

Freshly isolated PBMCs were stimulated with 250 U/ml of each type I IFN in R10 (3 x 10^6^ cells/sample) in a volume of 2 ml in a non-tissue culture treated 12 well plate and incubated for 24 h at 37°C. Cells were harvested by gentle pipetting, centrifuged at 300 x g for 5 min, resuspended in 1 ml R10 and the total cell number was determined. 0.5 x 10^6^ cells were transferred to a low-binding 1.5 ml tube and centrifuged at 300 x g for 5 min at 4°C. Cell pellets were resuspended in 100 μl of staining buffer (2% (v/v) BSA/0.01% (v/v) Tween-20 in cold RNase-free PBS + 10 μl Human TruStain FcX blocking solution (#422301, BioLegend)) and incubated for 10 min at 4°C. A mastermix of CITEseq antibodies (ADTs) was prepared consisting of 0.5 μg of each antibody and added to each sample together with 0.5 μg of unique Cell Hashing antibody (HTOs) (Supplementary Tables 16 and 17) and cells were stained for 30 min at 4°C. Cells were washed three times with 1 ml staining buffer, centrifuging at 350 x g for 5 min at 4°C. Cells were passed through a 35 μm cell strainer (Falcon) prior to the final wash step and were then resuspended in 200 μl staining buffer and counted. 1 x 10^5^ cells from each sample were pooled and passed over a 40 μm cell strainer (Falcon). After centrifugation at 350 x g for 5 min at 4°C, the cell pellet was resuspended in 200 ml cold RNase-free PBS to give a concentration of 1,500 cells/μl. Two lanes of a Chromium Single Cell Chip B (10X Genomics) were each superloaded with 30,000 cells, with a target of 15,000 single cells per lane.^65^ Generation of gel beads in emulsion (GEMs), GEM-reverse transcription, clean up and cDNA amplification were performed using a Chromium Single Cell 3′ Reagent Kit (v3), according to the manufacturer’s instructions, with the addition of 2 pmol each of ADT and HTO additive primers at the cDNA amplification step.^65^ During cDNA cleanup, the ADT- and HTO-containing supernatant fraction was separated from the cDNA fraction derived from cellular mRNAs using 0.6X SPRI beads (Beckman Coulter).

#### scRNAseq: Library generation and sequencing

The mRNA-derived cDNA library was prepared according to the standard Chromium Single Cell 3′ Reagent Kit (v3). ADTs and HTOs were purified using two 2X SPRI selection according to Stoeckius *et al*.^65^ and amplified using separate PCR reactions to generate the ADT and HTO libraries with ten cycles of amplification. PCR products were purified using 1.6X SPRI purification. Quality control of libraries was performed using a Qubit 3.0 dsDNA HS (high sensitivity) Assay Kit (Invitrogen) and BioAnalyzer High Sensitivity DNA Chip (Agilent). Libraries were diluted to 2 nM and pooled for sequencing in the following proportions: 80% cDNA, 10% ADT, 10% HTO. Libraries were sequenced twice on a NextSeq 500 with 150 bp paired-end reads (Illumina). Sequencing runs were pooled, yielding a total of 5 x 10^8^ reads for each 10X lane (1 x 10^9^ reads in total).

### Data analysis

#### Mass cytometry (CyTOF)

Data were normalised, randomised and concatenated using Helios CyTOF Software (v6.7) (Fluidigm). Compensation and debarcoding were performed in R (v3.5.1) using CATALYST (v1.5.3.23).^66^ FCS files were rewritten using the updatePanel function of cytofCore (v0.4). Intact, single, live, CD45+ cells were manually gated using FlowJo (v10.8) and exported as new FCS files. Data were transformed and cells clustered using FlowSOM as described in cytofWorkflow^67^ using CATALYST (v1.18.1) in R (4.1.3). Cell populations were manually identified by expression of known phenotyping markers and clusters merged and annotated.

#### Bulk RNAseq

Sequencing data were processed using a CGAT-core-based pipeline.^68^ Read quality was assessed using FastQC (v0.11.9) (bioinformatics.babraham.ac.uk/projects/fastqc). For the coding transcriptome, pseudoalignment was performed with Kallisto (v 0.46.1) using genome build GRCh38, with 34-49 million reads aligned per sample. For lncRNAs, pseudoalignment was performed with Kallisto (v0.46.1) using the full database (genome build GRCh38) from LNCipedia (v5.2).^69^ 10-14 million reads aligned per sample. Transcript abundances were imported into RStudio (v4.2.0) using tximport (v1.26.1)^70^ and summarised to gene level for the coding genes. Gene symbol annotations were added using AnnotationHub (v3.6.0). Differential expression analysis was performed using DESeq2 (v1.36.1)^71^ with a multi-factor design formula to account for the use of different donors. Non-expressed genes (genes with five counts or less in a sample) and genes that were expressed in two or fewer samples were excluded. Log fold change shrinkage (lfcShrink) for visualisation and gene ranking was performed using the shrinkage estimator “apeglm”. DEGs were defined as having an adjusted p value of < 0.05 and a log_2_ fold change of > 0.58 for up-regulated genes and < -0.58 for down-regulated genes. Gene ontology analysis was performed using goseq (v1.50.0).^72^ pheatmap (v1.0.12), ComplexHeatmap (v2.14.0), EnhancedVolcano (v1.16.0) and GraphPad Prism (v9.4.0) were used for data visualisation.

#### scRNAseq

BCL files were demultiplexed and converted to Fastq files using the mkfastq pipeline of Cell Ranger (v3.0.2) and bcl2fastq (v2.20.0.422) and read quality assessed using FastQC (v0.11.9). xRaw sequencing reads for the cDNA libraries were processed into count matrices using cellranger count (Cell Ranger v3.0.2) and aligned to Human reference 3.0.0 (GRCh38) (10X Genomics), which includes annotation of protein-coding genes, lncRNAs, antisense transcripts, immunoglobulin genes/pseudogenes and T cell receptor genes. Raw sequencing reads from the ADT and HTO libraries were processed into count matrices using CITE-Seq-Count (v1.4.2).^73^ The count matrices for cDNA, ADT and HTO libraries were imported into R (v4.1.3) and analysed using Seurat (v 4.1.1). Cells with < 200 or > 5000 genes or > 15% of mitochondrial reads were excluded. The HTO data was normalised using centred log-ratio (CLR) transformation and samples demultiplexed using the HTODemux function of Seurat. Count data for the RNA assay was normalised using sctransform v2, regressing out gene expression from ribosomal proteins and returning all genes. ADT data was normalised using CLR. Data from the two 10X lanes were merged and data scaled. The multimodal object was split into one object per sample and variable features normalised and identified using sctransform v2, using method “glGamPoi”. PrepSCTIntegration was performed on the six objects using all genes as anchor features and PCA performed. Integration anchors were identified and used to create an integrated data assay normalised using sctransform. Dimensionality reduction was performed using by Principal Component Analysis (PCA) using all features (genes) for the RNA assay and all features (antibodies) for the ADT assay. Weighted Nearest Neighbor (WNN) analysis^45^ was used to improve clustering by combining the RNA and ADT data. FindMultiModalNeighbors was performed using 30 dimensions for RNA and 18 for ADT and the data visualised using UMAP. Clusters were determined using using FindClusters using SLM algorithm with a resolution of 0.8 and cell types identified using expression of mRNA and protein markers.

Differential expression analysis was performed on the RNA assay which was first log normalised. The integrated object was subset by cell type and FindMarkers run for each type I IFN versus the unstimulated sample using a Wilcoxon Rank Sun test with a logfc threshold of 0.25 and genes that are detected in a minimum of 10% of cells. DEGs were defined as having an adjusted p value of < 0.05. Seurat (v4.1.1), Pheatmap (v1.0.12), ComplexHeatmap (v2.14.0), scCustomize (v1.1.1), eulerr (v7.0.0) and ggforce(v0.4.1.9000) were used for data visualisation.

## Supporting information

Key Resources Table

## Resource availability

### Lead contact

Further information and requests for resources and reagents should be directed to and will be fulfilled by the lead contact, Jan Rehwinkel.

### Materials availability

This study did not generate any unique reagents.

### Data and code availability

All data are available in the manuscript and associated supplementary files and at https://rehwinkellab.shinyapps.io/ifnresource/. Mass cytometry data generated during this study has been deposited at Flow Repository (FR-FCM-Z655) and is publicly available. The data used in Figure 1 was published previously.^63^ Sequencing data generated during this study has been deposited on ENA and is publicly available (PRJEB60774). All original code has been deposited at GitHub (https://github.com/rerigby/ifn-resource) and is publicly available. Any additional information required to reanalyse the data reported in this paper is available from the lead contact upon request.

## Acknowledgements

The authors would like to thank Clare Hardman and Yi-Ling Chen (MRC HIU) for assistance with phlebotomy, and Hayley Evans and David Fawkner-Corbett (MRC HIU) for technical advice. We would like to acknowledge Michalina Mazurczyk and Julia Menzies in the MRC WIMM Mass Cytometry Facility for providing technical expertise. The facility is supported by the MRC HIU core funded project reference MC_UU_00008 and the Oxford Single Cell Biology Consortium (OSCBC). We thank Neil Ashley in the MRC WIMM Single Cell Core Facility for his help with the CITE-seq experiments. The facility is supported by the MRC MHU (MC_UU_12009), the Oxford Single Cell Biology Consortium (MR/M00919X/1), the WT-ISSF (097813/Z/11/B#) and WIMM Strategic Alliance awards G0902418 and MC_UU_12025. We would like to acknowledge Tim Rostron in the Sequencing facility at the MRC WIMM for assistance with sequencing services. The facility is supported by the MRC HIU and by the EPA fund (CF268). This work was funded by the UK Medical Research Council [MRC core funding of the MRC Human Immunology Unit; J.R]. The funders had no role in study design, data collection and analysis, decision to publish, or preparation of the manuscript.

## Author contributions

Conceptualisation: R.E.R. and J.R.; Methodology: R.E.R.; Software: R.E.R. and K.R-A.; Validation: R.E.R. and J.R.; Formal analysis: R.E.R. and J.R.; Investigation: R.E.R; Resources: n/a; Data curation: R.E.R. and K.R-A.; Writing – Original Draft: R.E.R. and J.R.; Writing – Review & Editing: all authors; Visualisation: R.E.R., K.R-A. and J.R.; Supervision: J.R. and D.S.; Project administration: R.E.R. and J.R.; Funding acquisition: J.R.

## Supplementary Materials

**Supplementary Figures 1-26**

**Key Resources Table**

**Supplementary Tables 1-20**

<u>Supplementary Table 1:</u> Mass cytometry: Median expression of markers per cluster (IFN-α2a dose titration, 15 min stimulation), used in Figure 1

<u>Supplementary Table 2:</u> Mass cytometry: Number of cells per sample, by cluster (IFN-α2a dose titration, 15 min stimulation), used in Figure 1

<u>Supplementary Table 3:</u> Mass cytometry: Median expression of markers per cluster (all type I IFNs, 15 min stimulation), used in Figure 2

<u>Supplementary Table 4:</u> Mass cytometry: Number of cells per sample, by cluster (all type I IFNs, 15 min stimulation), used in Figure 2

<u>Supplementary Table 5:</u> Mass cytometry: Median expression of markers per cluster (IFN-α1, IFN-α2α, IFN-α10, IFN-β, IFN-ω, 90 min stimulation), used in Figure S4

<u>Supplementary Table 6:</u> Mass cytometry: Number of cells per sample, by cluster (IFN-α1, IFN-α2α, IFN-α10, IFN-β, IFN-ω, 90 min stimulation), used in Figure S4

<u>Supplementary Table 7:</u> Mass cytometry: Median expression of markers per cluster (IFN-α1, IFN-α2α, IFN-α10, IFN-β, IFN-ω, 24 h stimulation), used in Figure S5

<u>Supplementary Table 8:</u> Mass cytometry: Number of cells per sample, by cluster (IFN-α1, IFN-α2α, IFN-α10, IFN-β, IFN-ω, 24 h stimulation), used in Figure S5

<u>Supplementary Table 9:</u> Bulk RNAseq: Length scaled TPM (protein-coding genes)

<u>Supplementary Table 10:</u> Bulk RNAseq: Differential expression testing – all genes (protein-coding genes)

<u>Supplementary Table 11:</u> Bulk RNAseq: Significantly differentially expressed genes (protein-coding genes)

<u>Supplementary Table 12:</u> Curated list of ISGs, used in Figure 3 and Figure S12

<u>Supplementary Table 13:</u> Bulk RNAseq: Length scaled TPM (lncRNAs)

<u>Supplementary Table 14:</u> Bulk RNAseq: Differential expression testing – all lncRNAs (lncRNAs)

<u>Supplementary Table 15:</u> Bulk RNAseq: Significantly differentially expressed genes (lncRNAs)

<u>Supplementary Table 16:</u> CITE-seq: HTO sequences

<u>Supplementary Table 17:</u> CITE-seq: ADT sequences

<u>Supplementary Table 18:</u> CITE-seq: Differentially expressed genes (IFN-α1 v unstim)

<u>Supplementary Table 19:</u> CITE-seq: Differentially expressed genes (IFN-α2a v unstim)

<u>Supplementary Table 20:</u> CITE-seq: Differentially expressed genes (IFN-α10 v unstim)

<u>Supplementary Table 21:</u> CITE-seq: Differentially expressed genes (IFN-β v unstim)

<u>Supplementary Table 22:</u> CITE-seq: Differentially expressed genes (IFN-ω v unstim)

**Figure S1 (Related to Figure 1).**
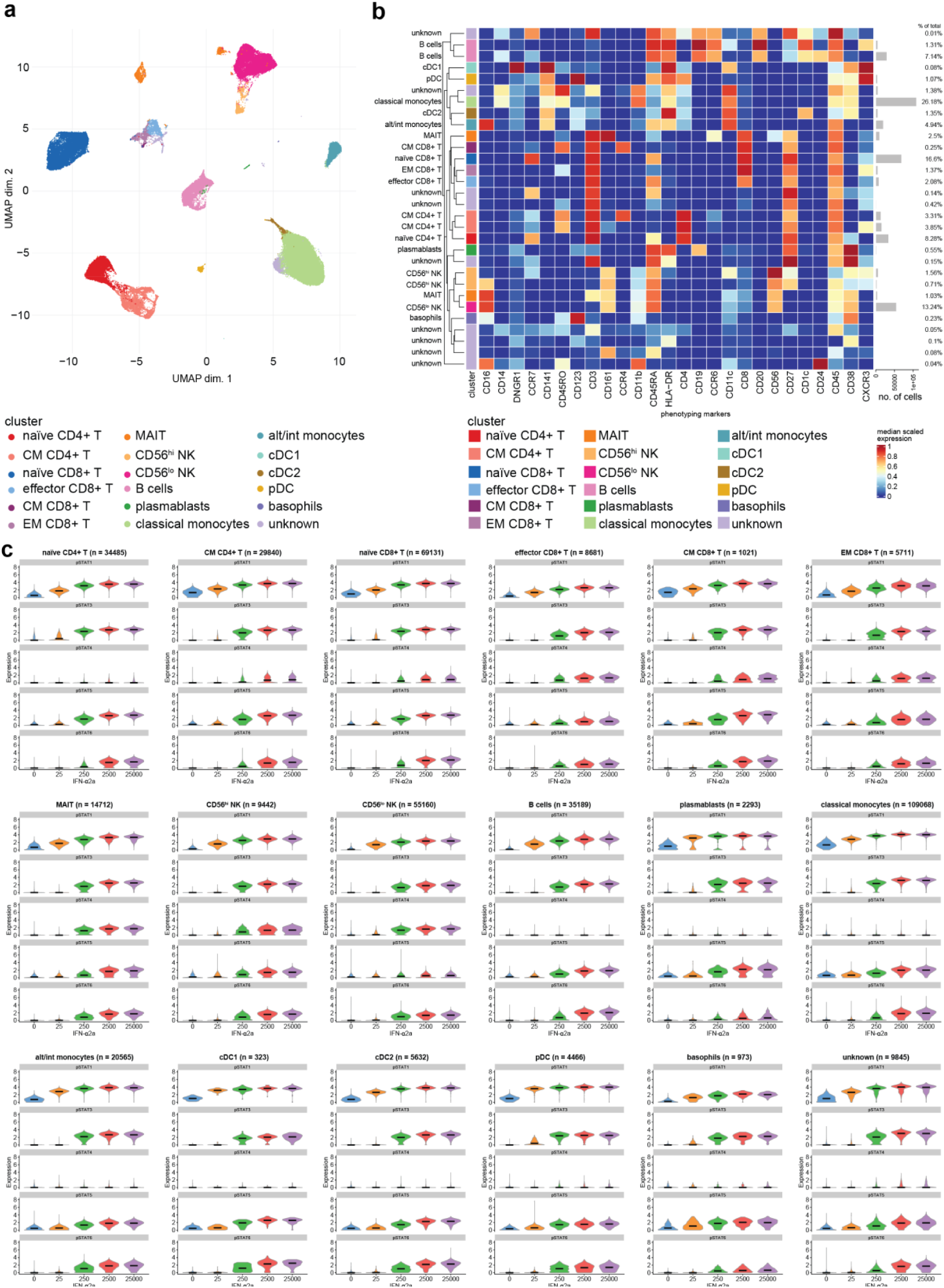
Phosphorylation of STAT proteins in response to IFN-α2a at 15 minutes. (a, b) UMAP (a) and heatmap (b) plots showing clustering of PBMCs and identification of different cell types based on expression of phenotyping markers. The percentage of cells per cluster is shown in the histogram on the right-hand side of (b). (c) Violin plots showing expression of each pSTAT in the indicated cell types in response to increasing concentrations of IFN-α2a.

**Figure S2 (Related to Figure 1).**
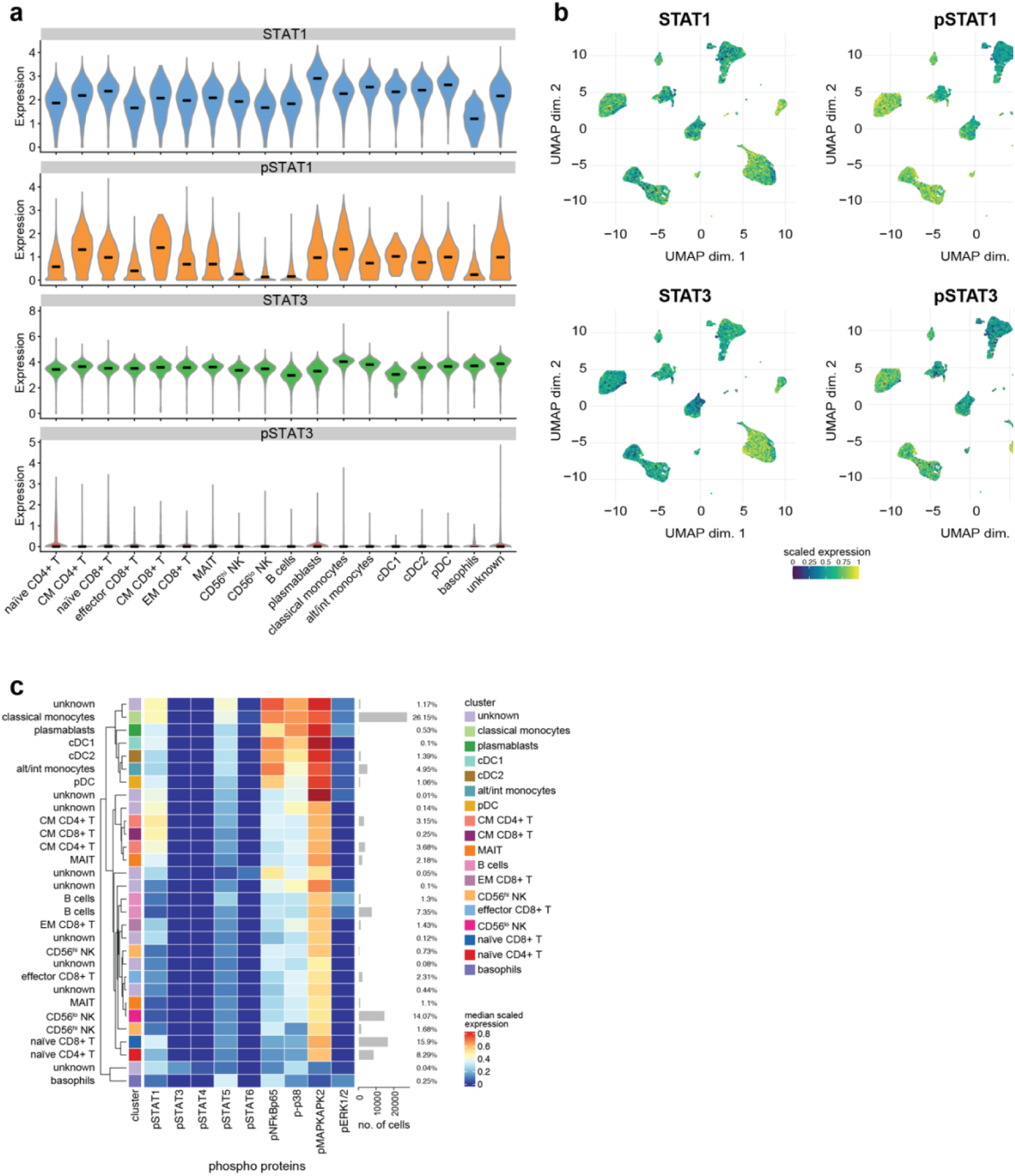
Baseline expression of STAT1, pSTAT1, STAT3 and pSTAT3 in different cell types. (a, b) Violin (a) and UMAP (b) plots showing the expression of STAT1, pSTAT1, STAT3 and pSTAT3 in unstimulated PBMCs, clustered as shown in Figure S1a. (c) Heatmap showing expression of phosphorylated proteins in unstimulated cells.

**Figure S3 Related to Figure 2).**
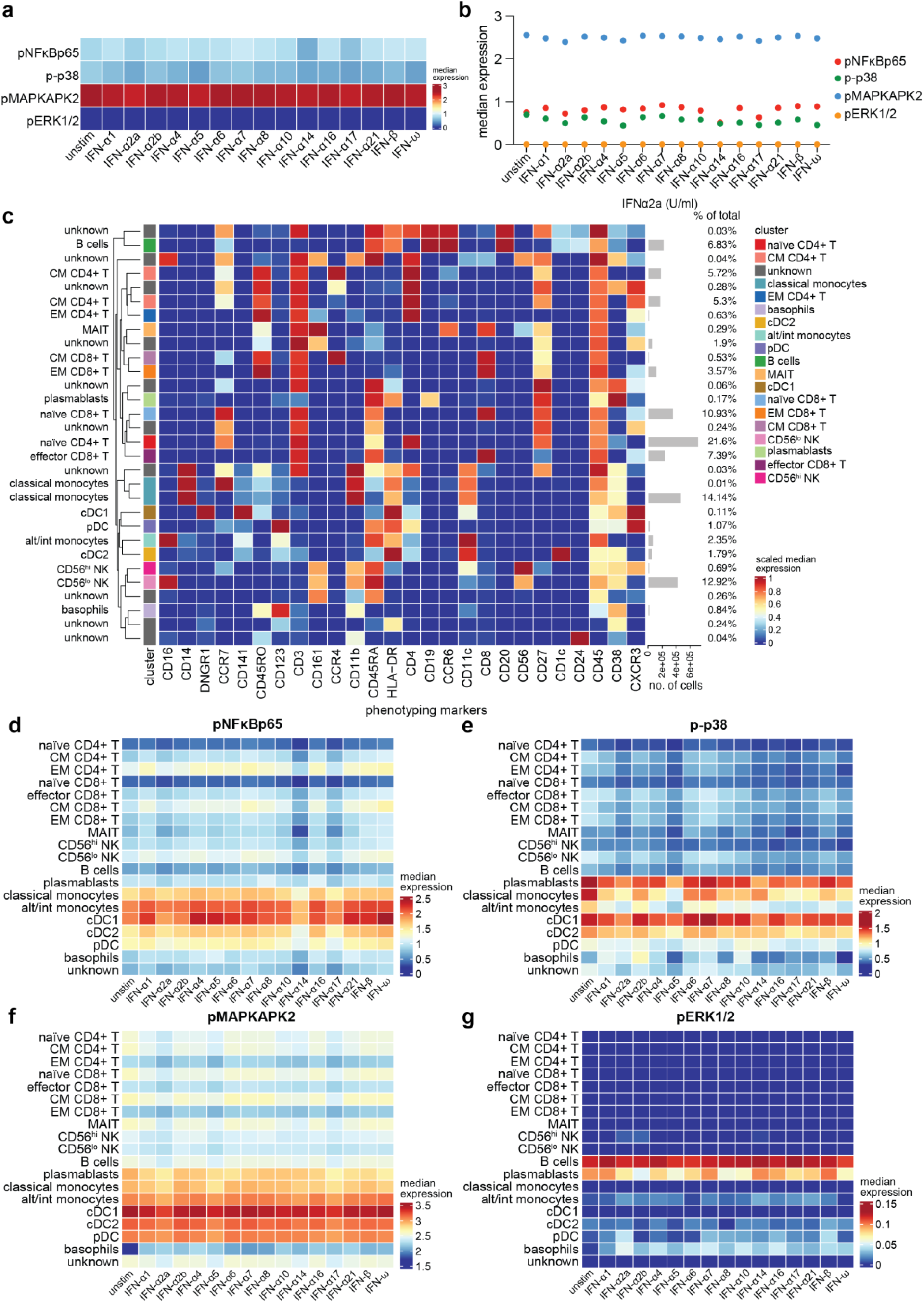
(Phosphorylation of signalling proteins in response to type I IFN stimulation at 15 minutes. (a) Median expression of pNFκBp65, p-p38, pMAPKAPK2 and pERK1/2 in PBMCs in response to treatment with 2,500 U/ml of each type I IFN for 15 minutes. (b) Depiction of the data shown in (a) as a dot plot. (c) Heatmap showing expression of the 26 phenotyping markers used for cell type identification as shown in Figure 2c. The percentage of cells per cluster is shown in the histogram on the right-hand side. (d-g) Heatmaps showing median expression of the indicated phosphoproteins in each cell type in response to treatment with different type I IFNs.

**Figure S4 (Related to Figure 2).**
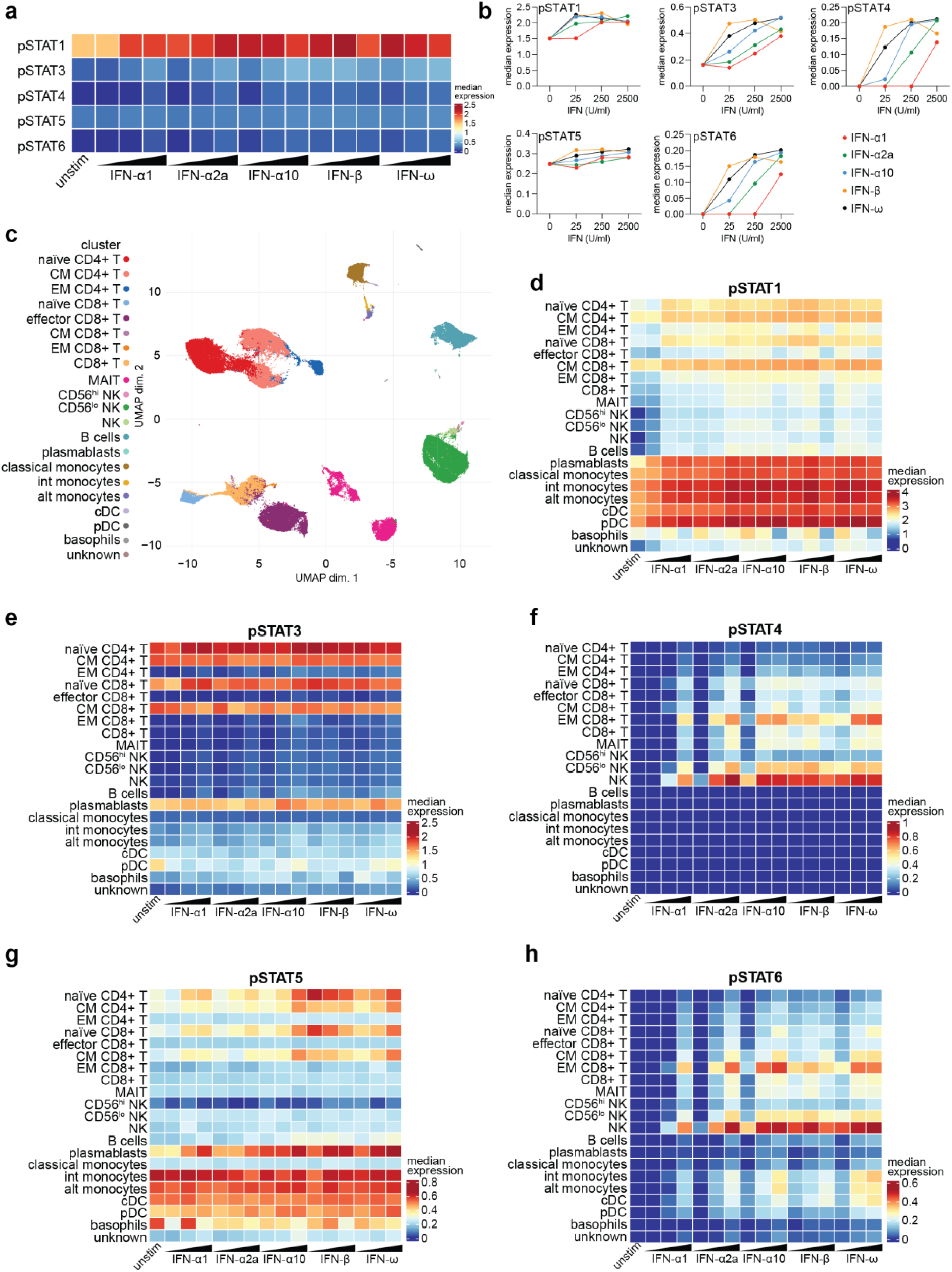
Phosphorylation of STAT proteins in response to stimulation with type I IFNs at 90 minutes. (a) Median expression of pSTATs in PBMCs in response to treatment with 25, 250 and 2,500 U/ml of the indicated type I IFNs for 90 minutes. (b) Depiction of the data shown in (a) as line plots. (c) UMAP plot showing clustering of PBMCs and identification of different cell types based on expression of the phenotyping markers. (d-h) Heatmaps showing median expression of pSTATs in each cell type in response to treatment with different type I IFNs.

**Figure S5 (Related to Figure 2).**
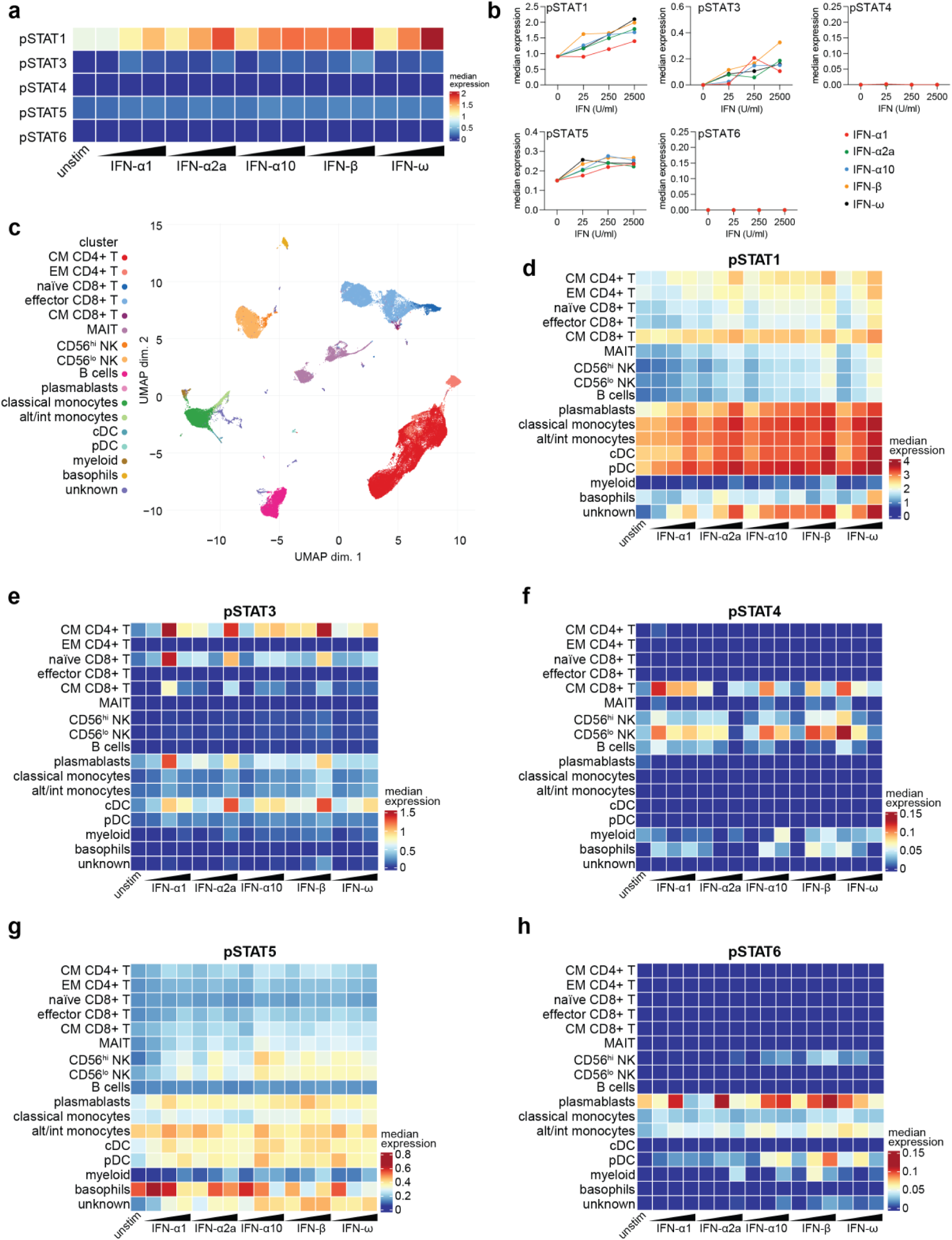
Phosphorylation of STAT proteins in response to stimulation with type I IFNs at 24 hours. (a) Median expression of pSTATs in PBMCs in response to treatment with 25, 250 and 2,500 U/ml of the indicated type I IFNs for 24 hours. (b) Depiction of the data shown in (a) as line plots. (c) UMAP plot showing clustering of PBMCs and identification of different cell types based on expression of the phenotyping markers. (d-h) Heatmaps showing median expression of pSTATs in each cell type in response to treatment with different type I IFNs.

**Figure S6 (Related to Figure 2).**
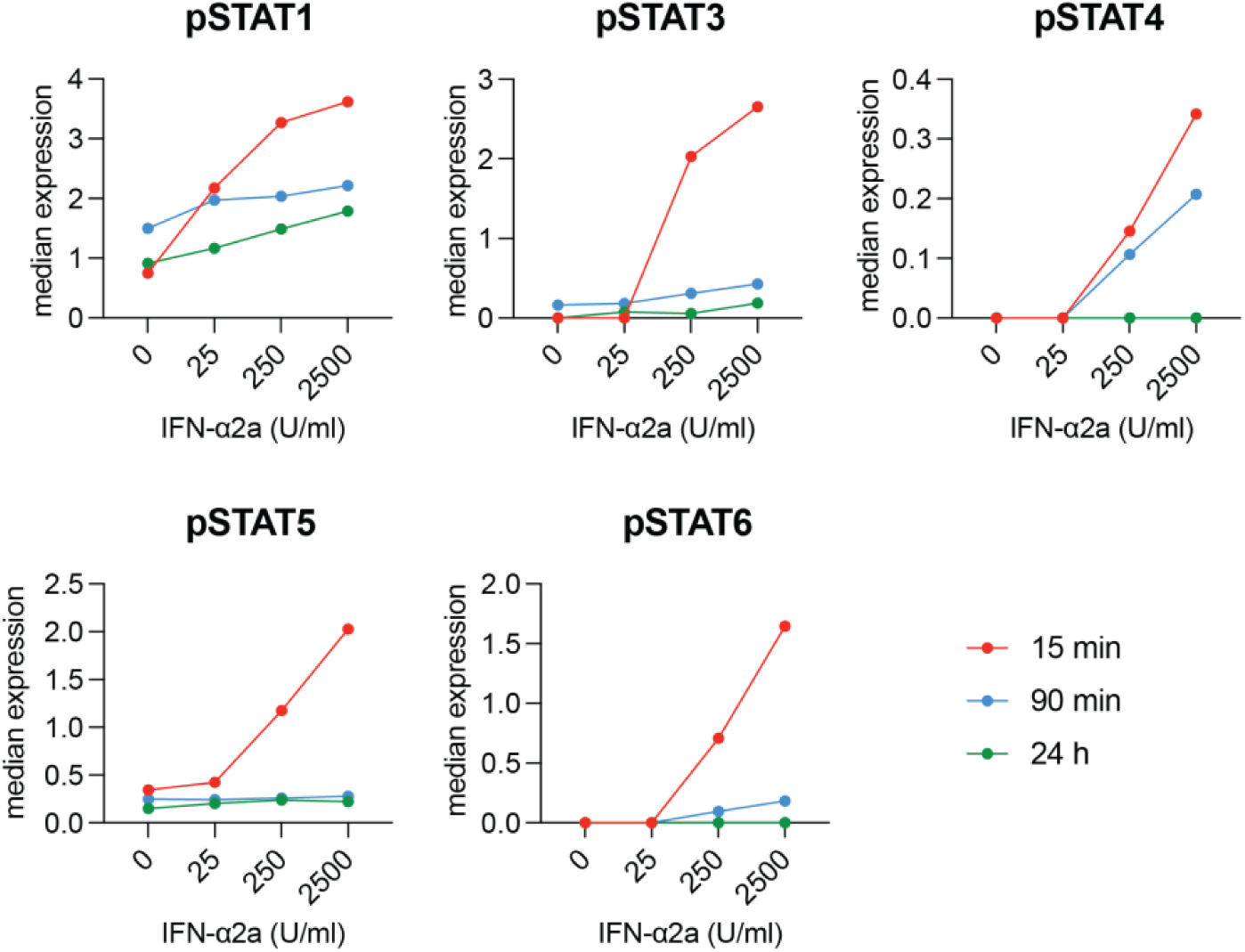
Phosphorylation of STAT proteins in response to IFN-α2a at different timepoints. Median expression of pSTATs in total PBMCs after stimulation with IFN-α2a for 15 minutes, 90 minutes or 24 hours. The data were pooled from Figures 1c, S4b and S5b without further normalisation.

**Figure S7 (Related to Figure 3).**
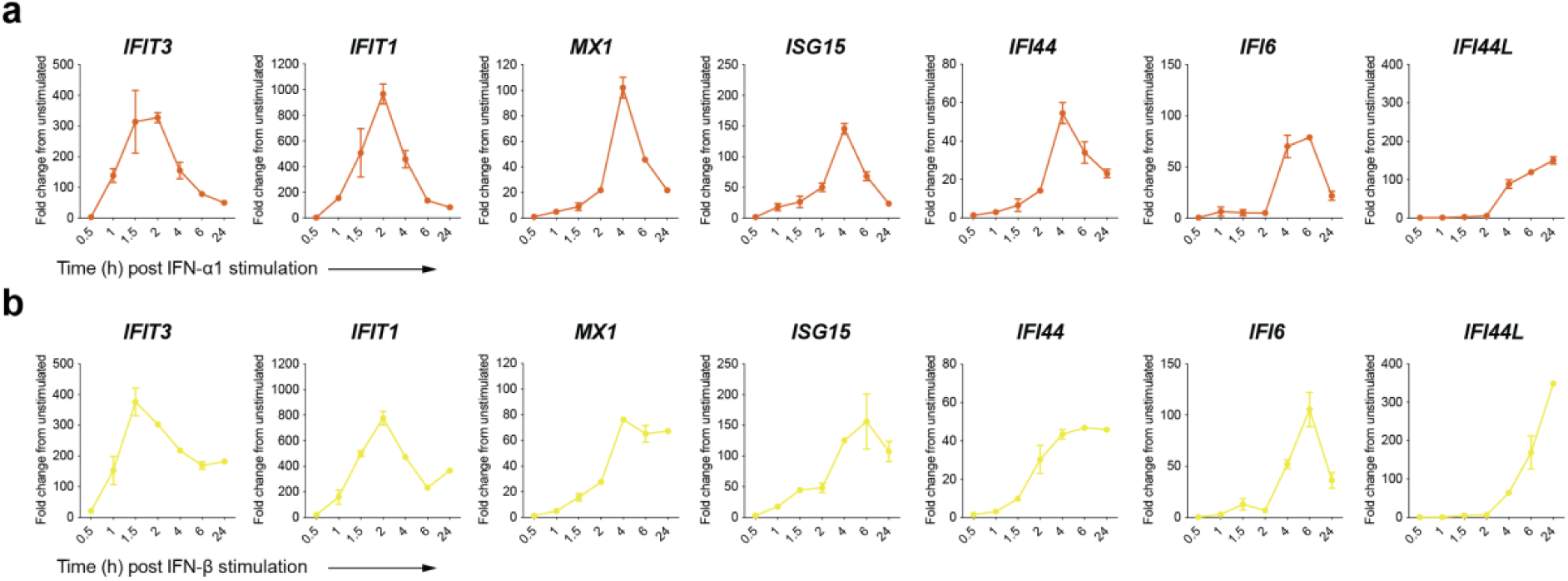
Kinetics of ISG expression in response to stimulation with IFN-α1 or IFN-β. (a, b) PBMCs were stimulated with 250 U/ml of IFN-α1 (a) or IFN-β (b) or left unstimulated for the indicated periods of time. RNA was extracted and RT-qPCR for the indicated ISGs was performed. Data were normalised to expression of the housekeeping gene *HPRT* and are shown as fold change relative to unstimulated cells harvested at the same timepoint. Data are from PBMCs from one donor and error bars show range of duplicate stimulated wells.

**Figure S8 (Related to Figure 3).**
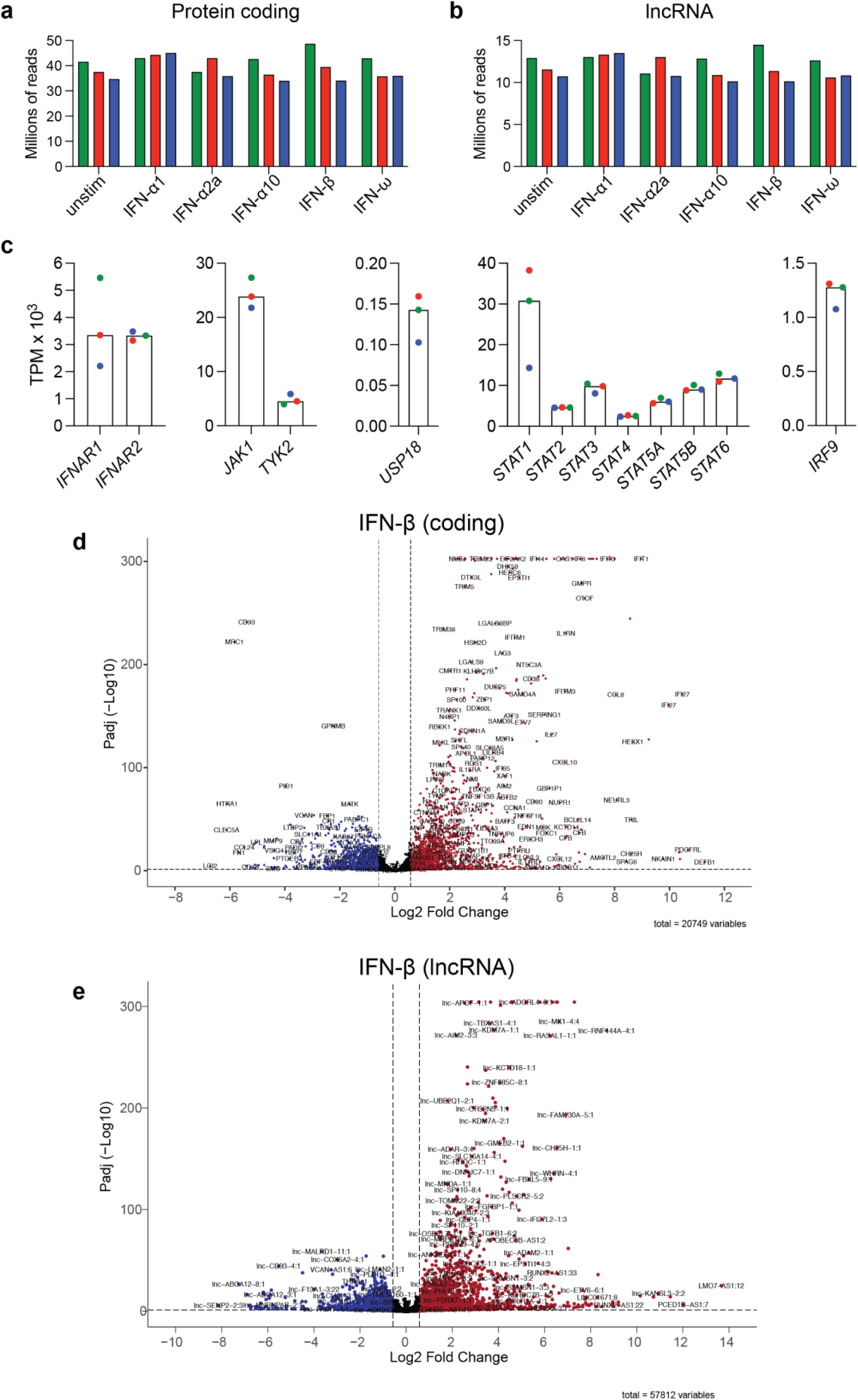
Differentially expressed protein-coding genes and lncRNAs in type I IFN-stimulated samples. (a, b) Number of reads pdeudoaligning with the protein-coding human transcriptome (a) or lncRNA database (b). (c) Quantification of expression of the indicated genes. (d,e) Labelled volcano plots for IFN-β-stimulated samples from Figure 3a and 3b. In (a-c), colours indicate individual PBMC donors.

**Figure S9 (Related to Figure 3).**
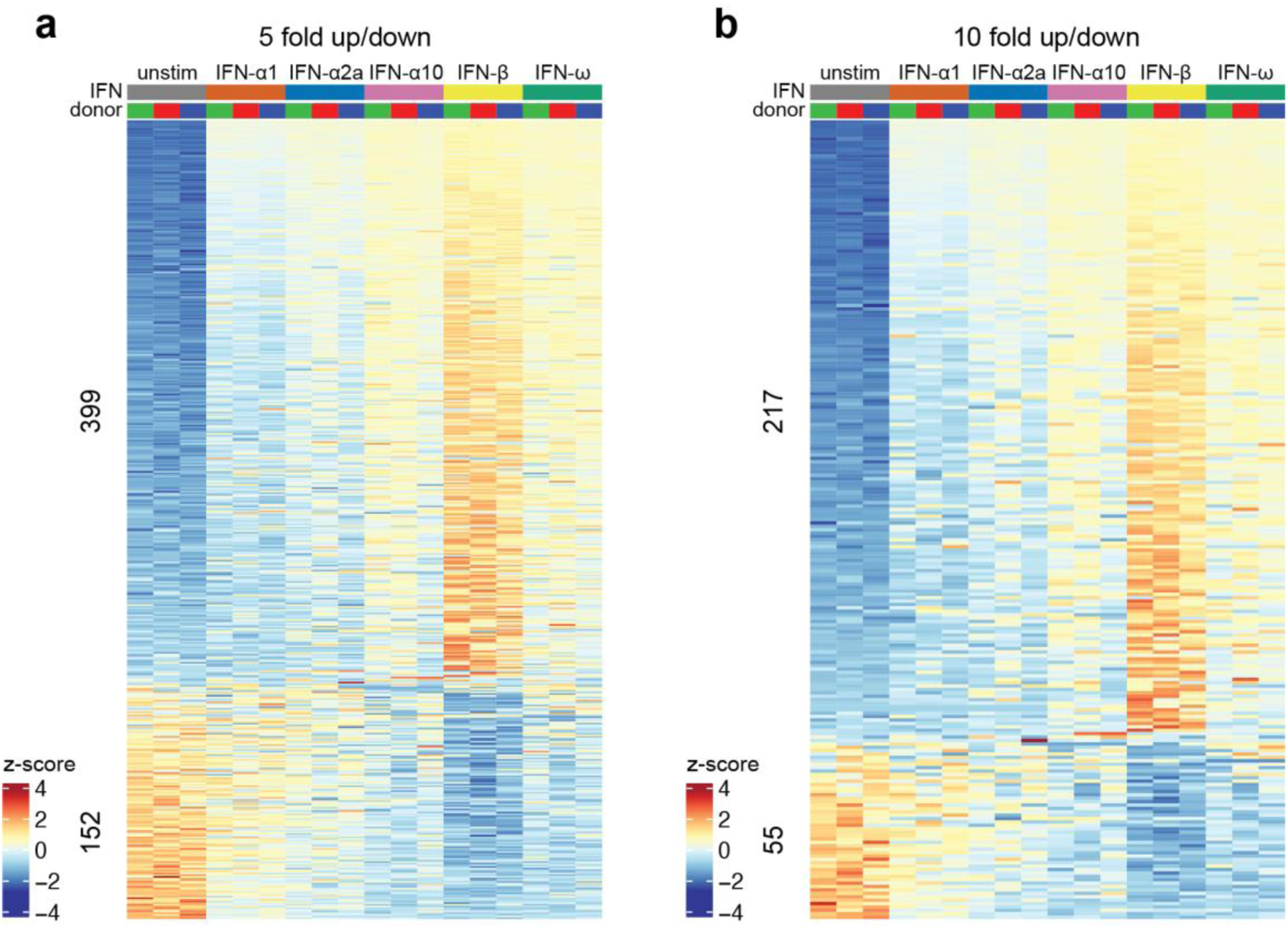
Number of differentially expressed genes using increased fold change stringency filters. (a,b) Heatmaps of genes differentially expressed in response to stimulation with at least one type I IFN (padj < 0.05) using 5-fold (a) and 10-fold (b) thresholds. Genes are ranked by z-score of all type I IFN-stimulated samples.

**Figure S10 (Related to Figure 3).**
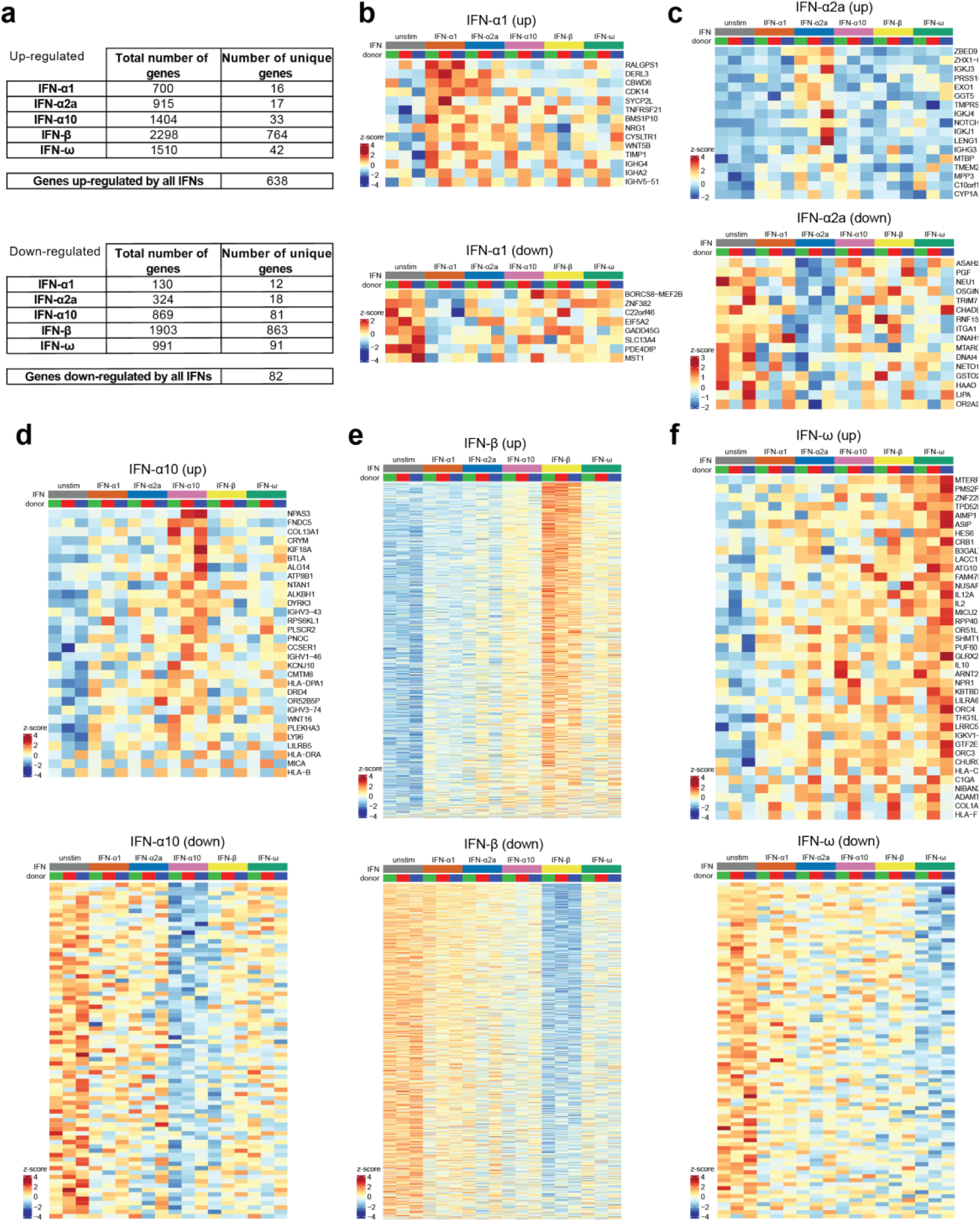
Genes differentially expressed in response to one type I IFN subtype only. (a) Summary table of the number of differentially expressed protein-coding genes in response to stimulation with each type I IFN subtype (padj < 0.05, fold change > 1.5 or < -1.5). (b-f) Heatmaps for genes uniquely up-or down-regulated by the indicated type I IFN subtype compared to unstimulated cells. Row names were omitted for clarity from heatmaps with > 50 rows for visualisation purposes. Genes were ranked by z-score for the indicated type I IFN-stimulated samples.

**Figure S11 (Related to Figure 3).**
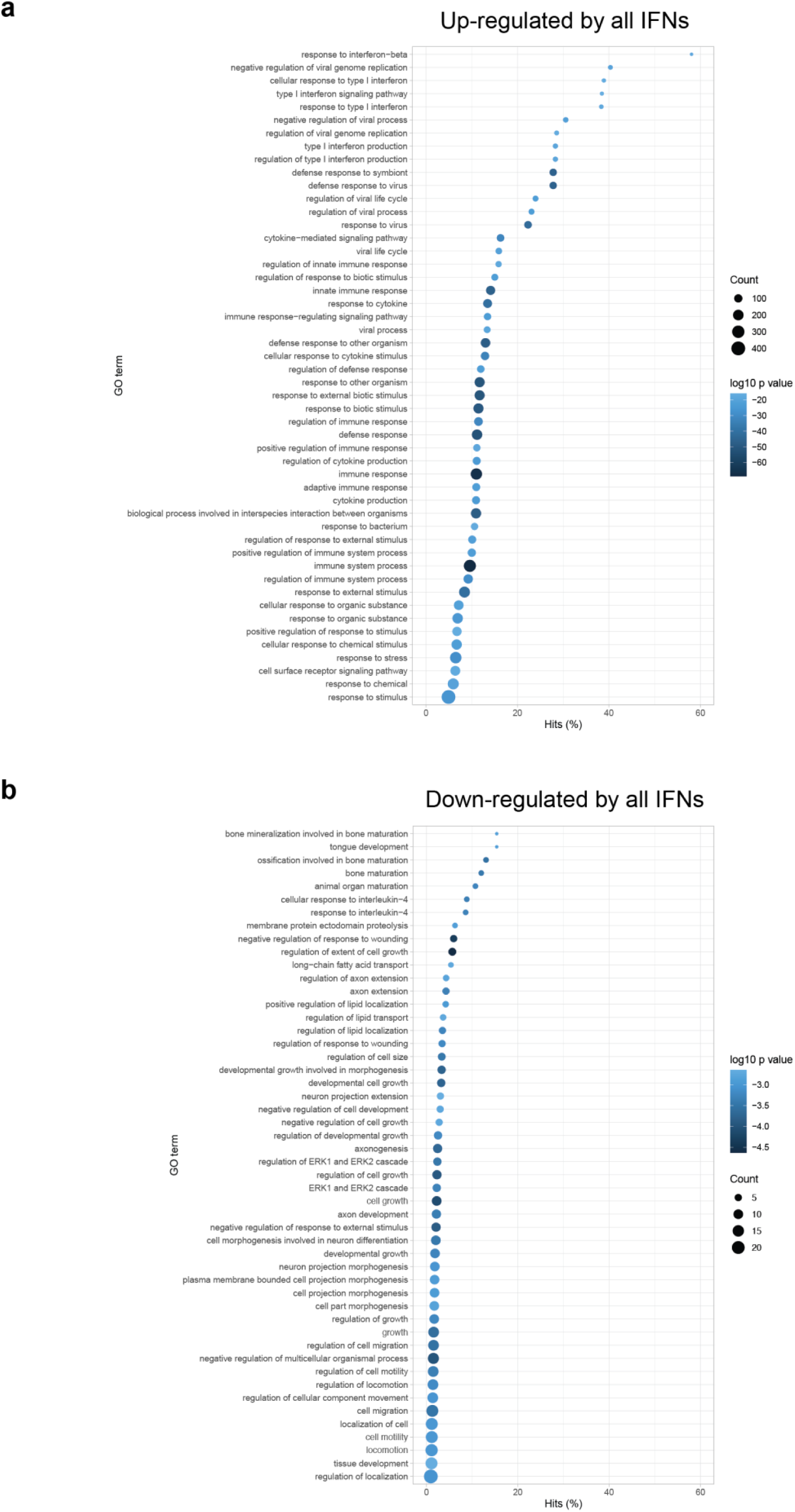
Gene Ontology of genes significantly differentially expressed by all type I IFNs compared to unstimulated PBMCs. (a,b) Gene Ontology (GO) analysis was performed using the lists of genes significantly up-(padj < 0.05, fold change > 1.5) or down-(padj < 0.05, fold change < -1.5) regulated by all type I IFNs using GOSeq, accounting for gene length. The Top 50 Biological Process GO terms are shown for upregulated (a) and downregulated (b) genes, sorted according to the percentage of hits for each term from the genes in each list. The number of genes and p-value for each GO term are indicated.

**Figure S12 (Related to Figure 3).**
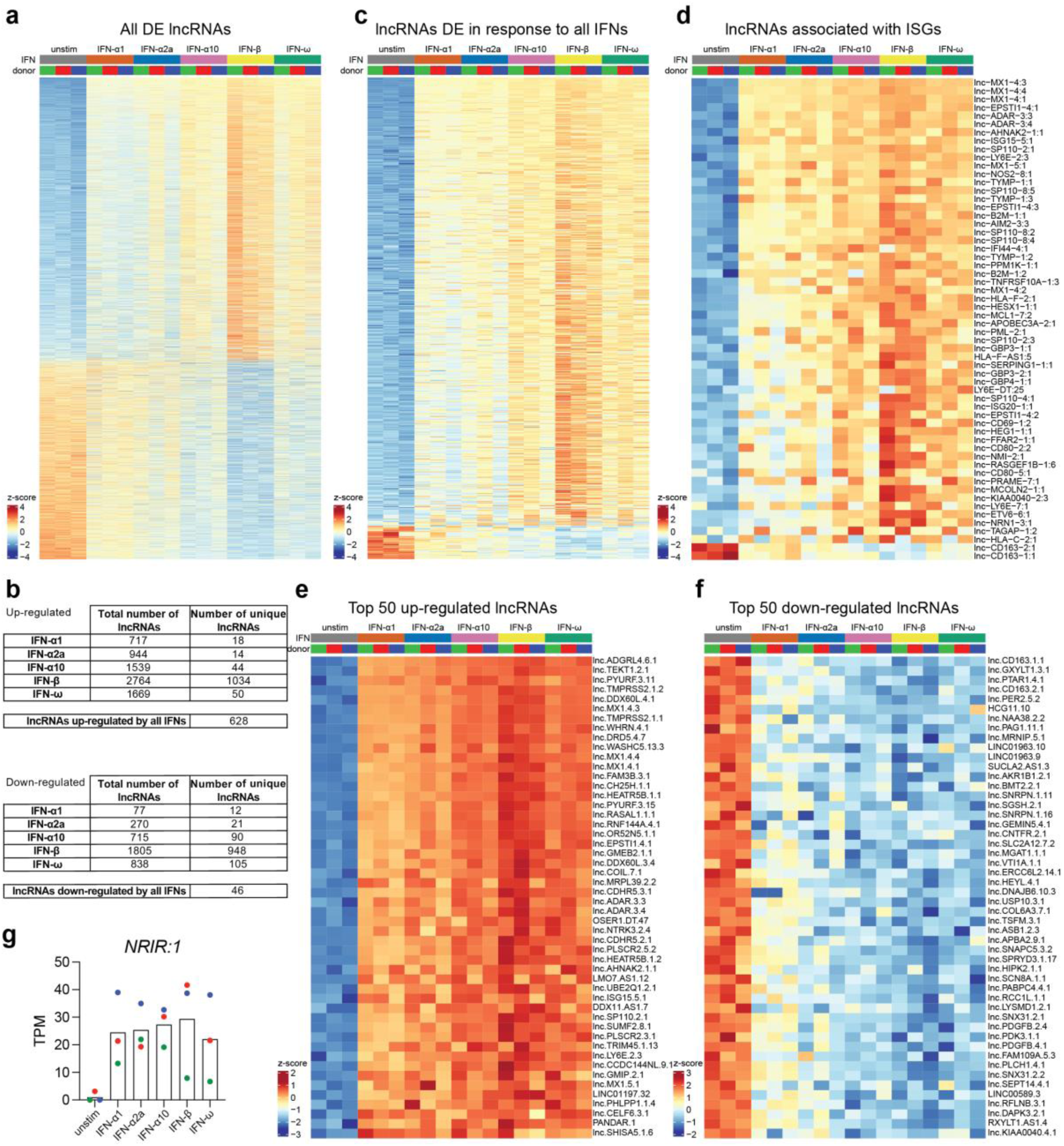
Differential expression of lncRNAs following stimulation with type I IFN compared to unstimulated PBMCs. (a) Heatmap of all lncRNAs significantly differentially expressed (padj < 0.05, fold change > 1.5 or < -1.5) in response to at least one type I IFN, compared to unstimulated PBMCs (2,931 up-regulated lncRNAs and 2,053 down-regulated lncRNAs). (b) Number of lncRNAs differentially expressed in response to each different type I IFN subtype. (c) lncRNAs differentially expressed in response to all type I IFN subtypes. (d) lncRNAs shown in (c) that are associated with ISGs (n = 56 up-regulated and 2 down-regulated). (e,f) Top 50 up-regulated (e) and down-regulated (f) lncRNAs. (g) Quantification of lncRNA *NRIR*. Colours indicate individual PBMC donors. In (a) and (c-f), lncRNAs are ranked by z-score of all type I IFN-stimulated samples.

**Figure S13 (Related to Figure 4).**
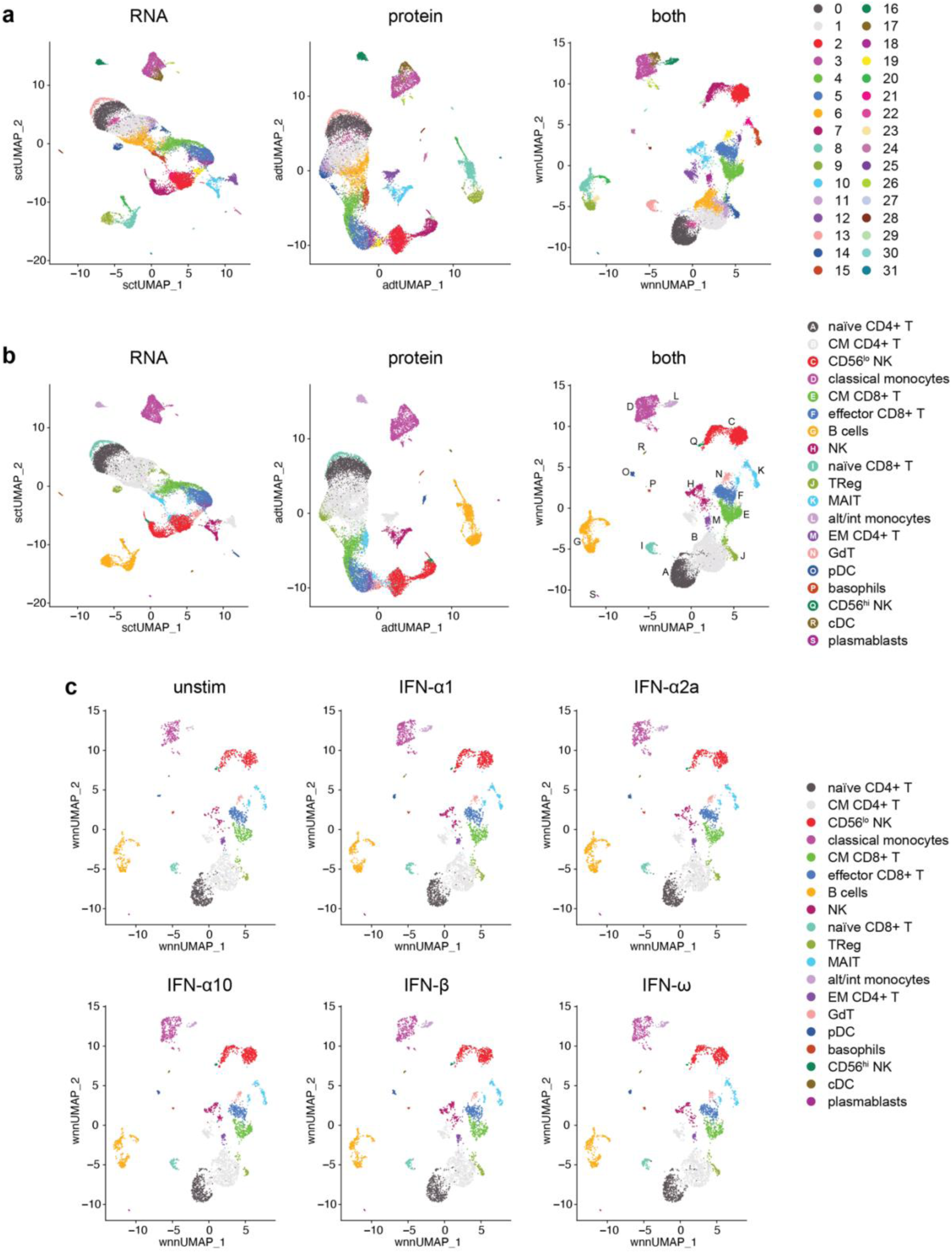
CITE-seq of PBMCs stimulated with type I IFNs. (a) UMAP plots showing the 32 cell clusters identified following clustering by RNA expression, protein expression or a weighted combination of both (cells from all samples combined). (b) Identification of different cell types based on merging of the clusters in (a). (c) Separation of the WNN UMAP by sample.

**Figure S14 (Related to Figure 4).**
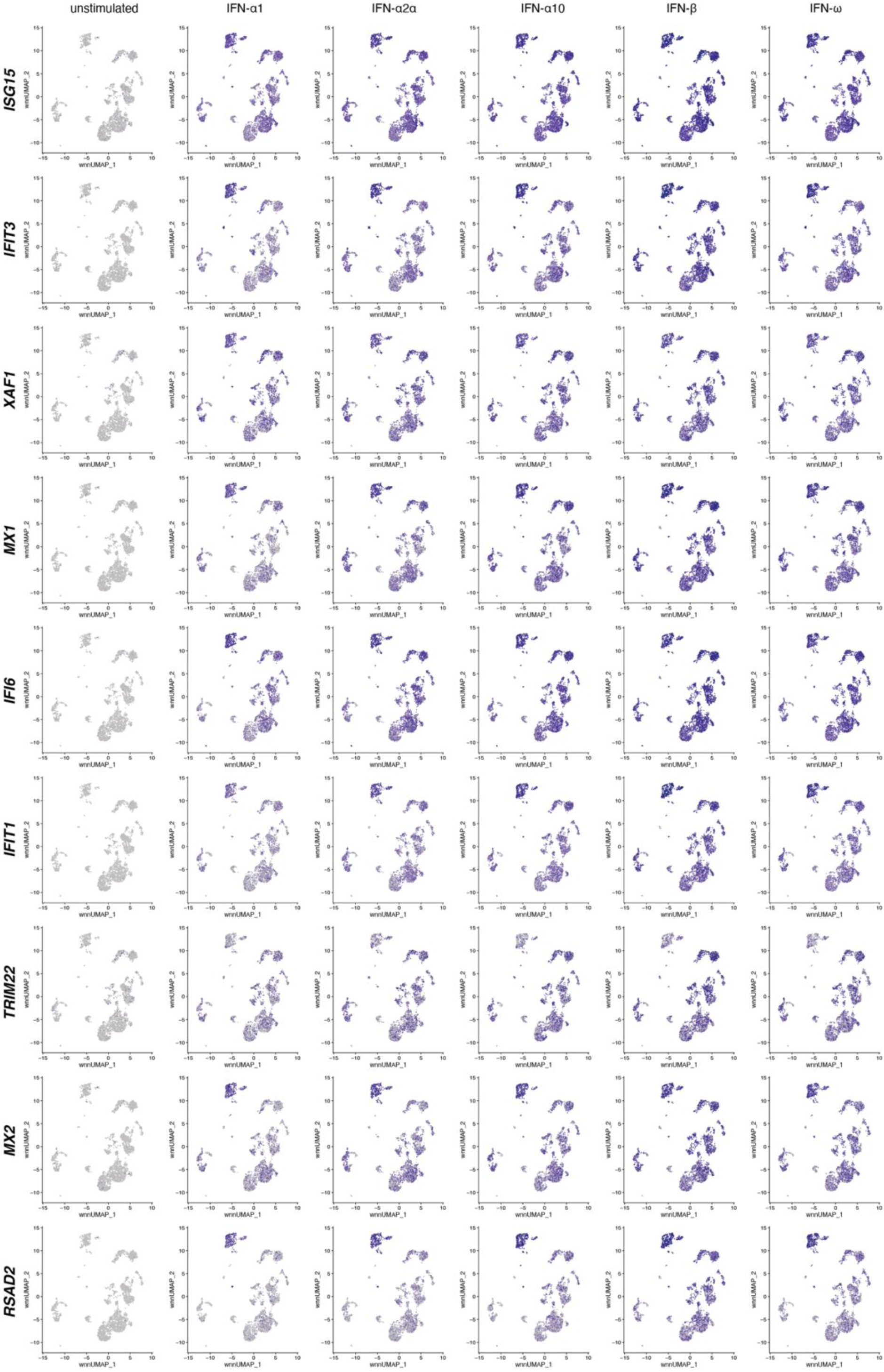
Core ISGs. UMAP plots showing the expression of the ten core ISGs across all samples.

**Figure S15 (Related to Figure 5).**
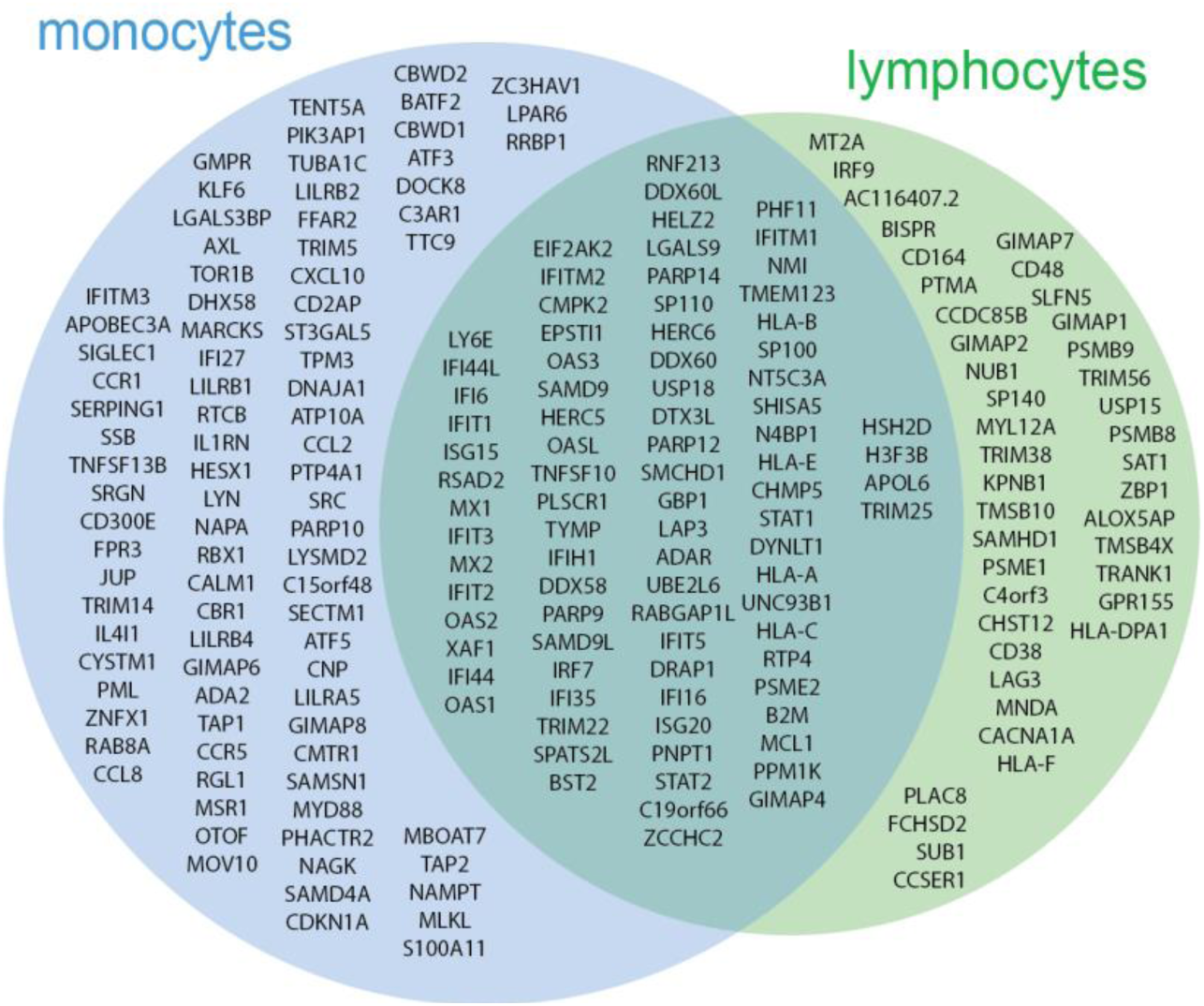
Monocyte and lymphocyte-specific ISGs. Venn diagram from Figure 5a showing the names of the genes in each segment.

**Figure S16 (Related to Figure 5).**
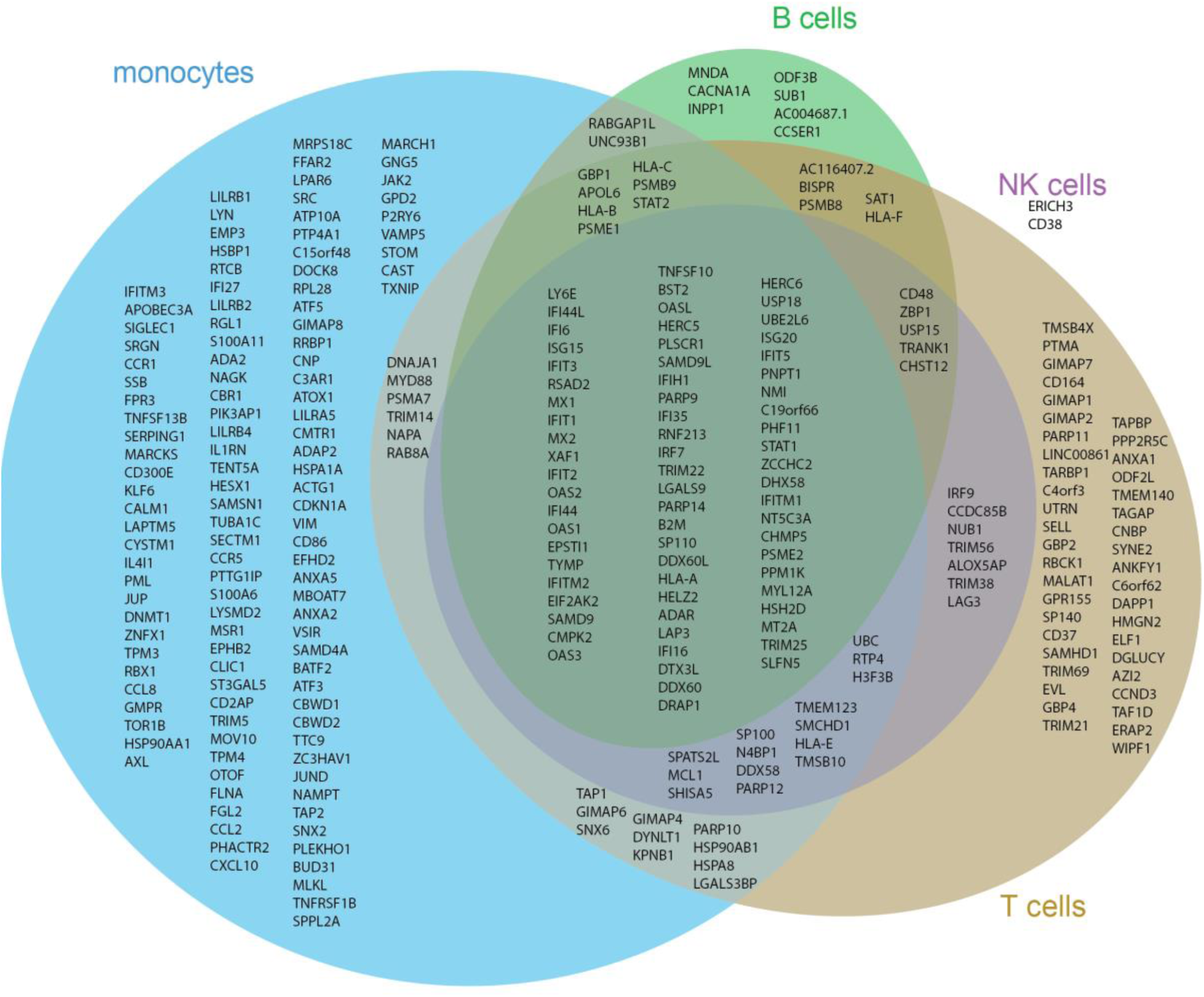
Monocyte and lymphocyte-specific ISGs. Venn diagrams showing cell type-specific ISGs as in Figure S15, further dividing lymphocytes into B cells, T cells and NK cells.

**Figure S17 (Related to Figure 5).**
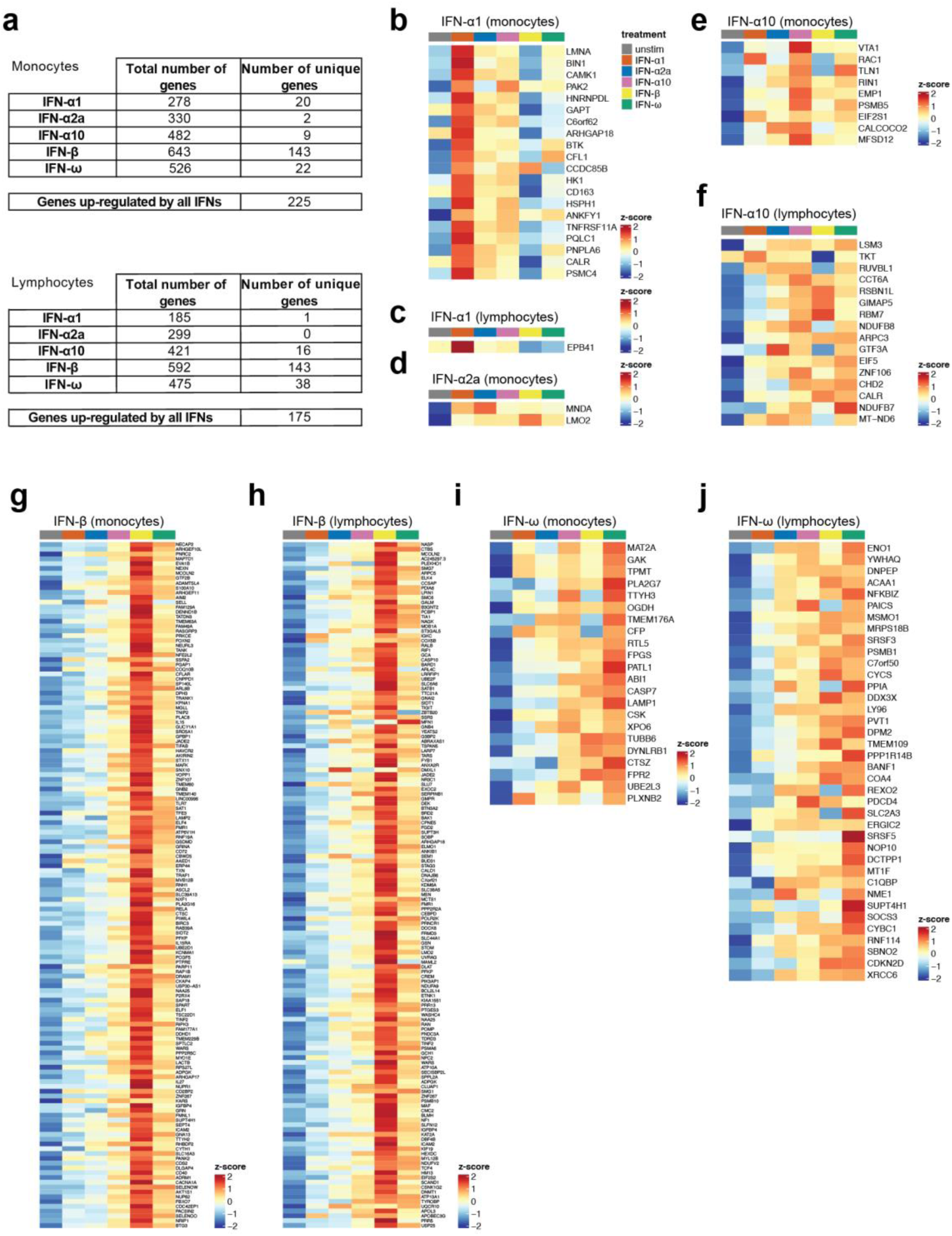
Genes up-regulated in response to only one type I IFN subtype only in monocytes or lymphocytes. (a) Summary tables of the number of differentially expressed genes in response to stimulation with each type I IFN subtype in monocytes (classical monocytes, alternative/intermediate monocytes) and lymphocytes (central memory CD4+ T cells, naïve CD4+ T cells, CD56^lo^ NK cells, B cells, central memory CD8+ T cells, effector CD8+ T cells, MAIT cells, other NK cells, naïve CD8+ T cells, regulatory T cells, effector memory CD4+ T cells and γδ-T cells). (b-j) Heatmaps for genes uniquely up-regulated by the indicated type I IFN subtype compared to unstimulated PBMCs (padj < 0.05, log_2_ fold change > 0.25) plotted for myeloid cells (all monocytes, cDCs, basophils) or lymphocytes (all T cells, B cells, NK cells, pDCs). Genes were ranked by z-score for the indicated type I IFN-stimulated samples.

**Figure S18 (Related to Figure 6).**
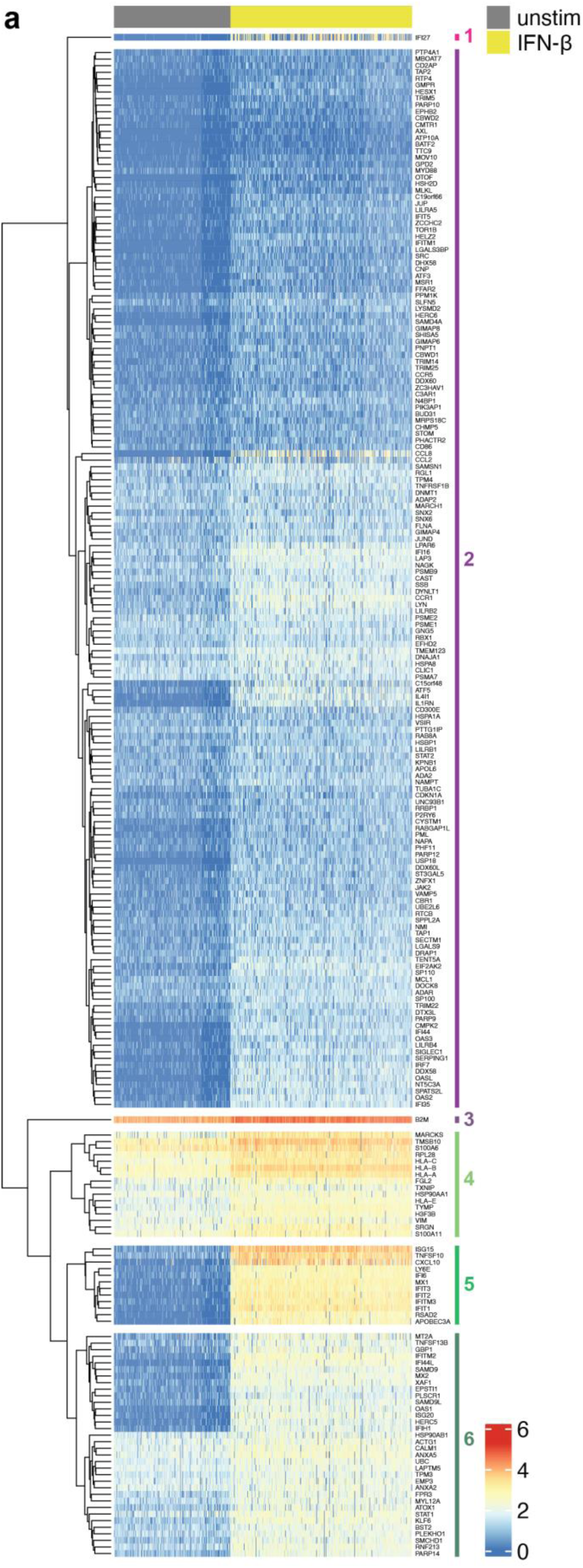
Response of classical monocytes to type I IFNs. Enlargement of the heatmap in Figure 6a with annotated rows (for online viewing).

**Figure S19 (Related to Figure 6).**
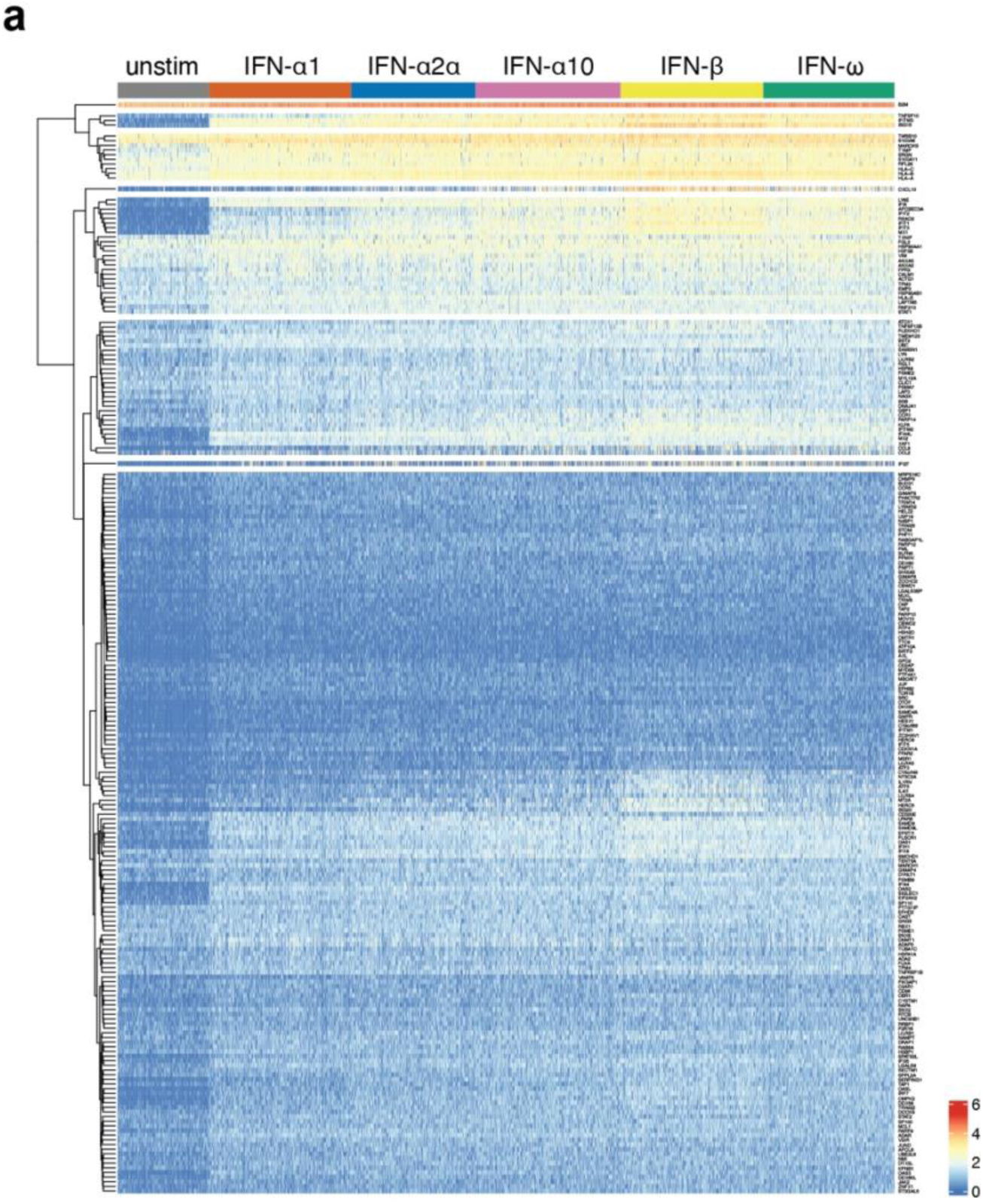
Response of classical monocytes to type I IFNs. Heatmap showing expression of the 225 genes significantly up-regulated in response to all tested type I IFNs in classical monocytes, in unstimulated and stimulated cells. Each row represents a gene and each column a cell.

**Figure S20 (Related to Figure 6).**
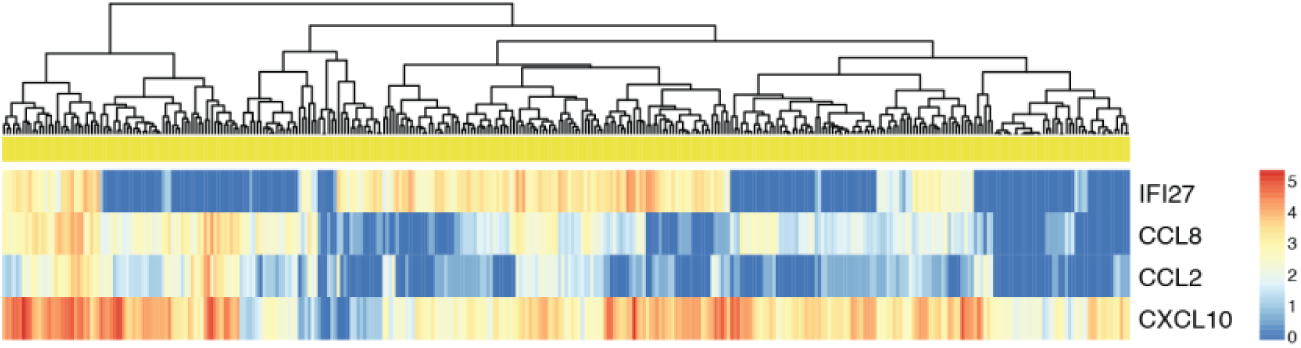
Expression of *IFI27*, *CCL8*, *CCL2* and *CXCL10* in individual cells. Expression of the indicated genes is shown for IFN-β-stimulated classical monocytes. Heatmaps are clustered by columns which represent individual cells.

**Figure S21 (Related to Figure 6).**
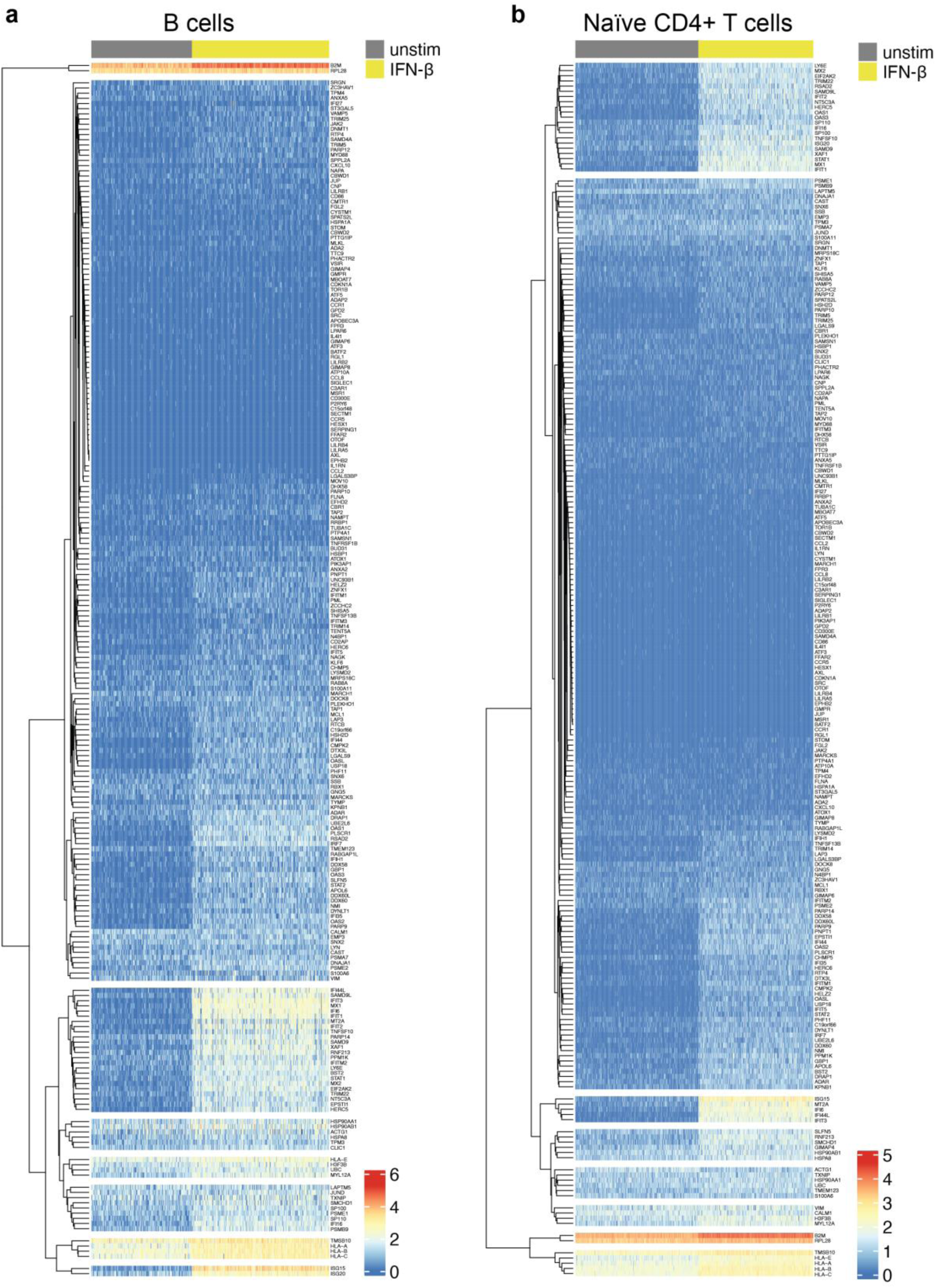
Expression of genes up-regulated in classical monocytes in other cell types. Heatmaps showing expression of the 225 genes that were significantly up-regulated by all tested type I IFNs in classical monocytes, in unstimulated and IFN-β-treated B cells and naïve CD4+ T cells. Gene names are provided for online viewing.

**Figure S22 (Related to Figure 6).**
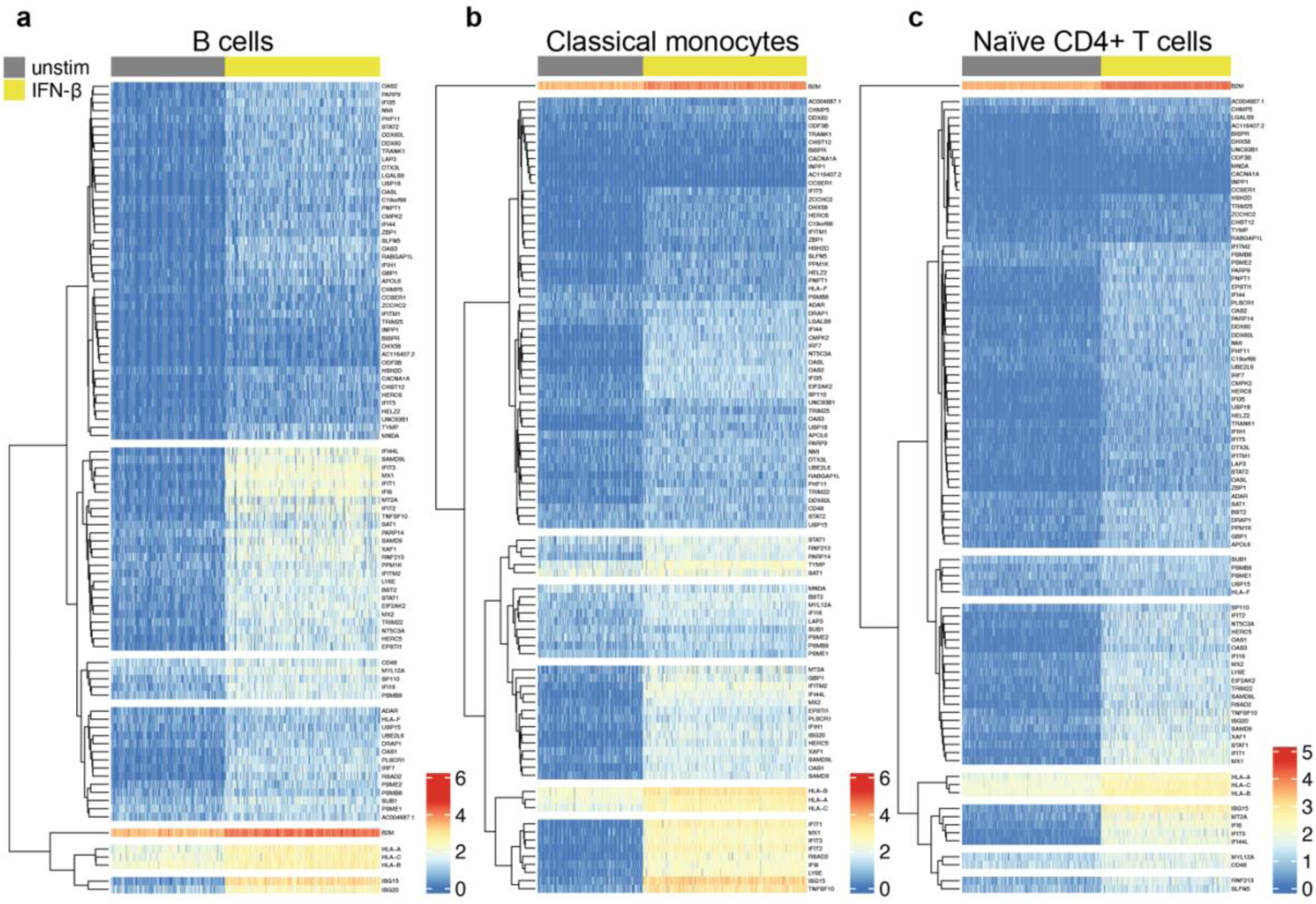
Response of B cells to type I IFNs. (a-c) Heatmaps showing expression of the 94 genes significantly up-regulated by all tested type I IFNs in B cells in unstimulated and IFN-β-treated cells for B cells (a), classical monocytes (b) and naïve CD4+ T cells (c). Gene names are provided for online viewing.

**Figure S23 (Related to Figure 6).**
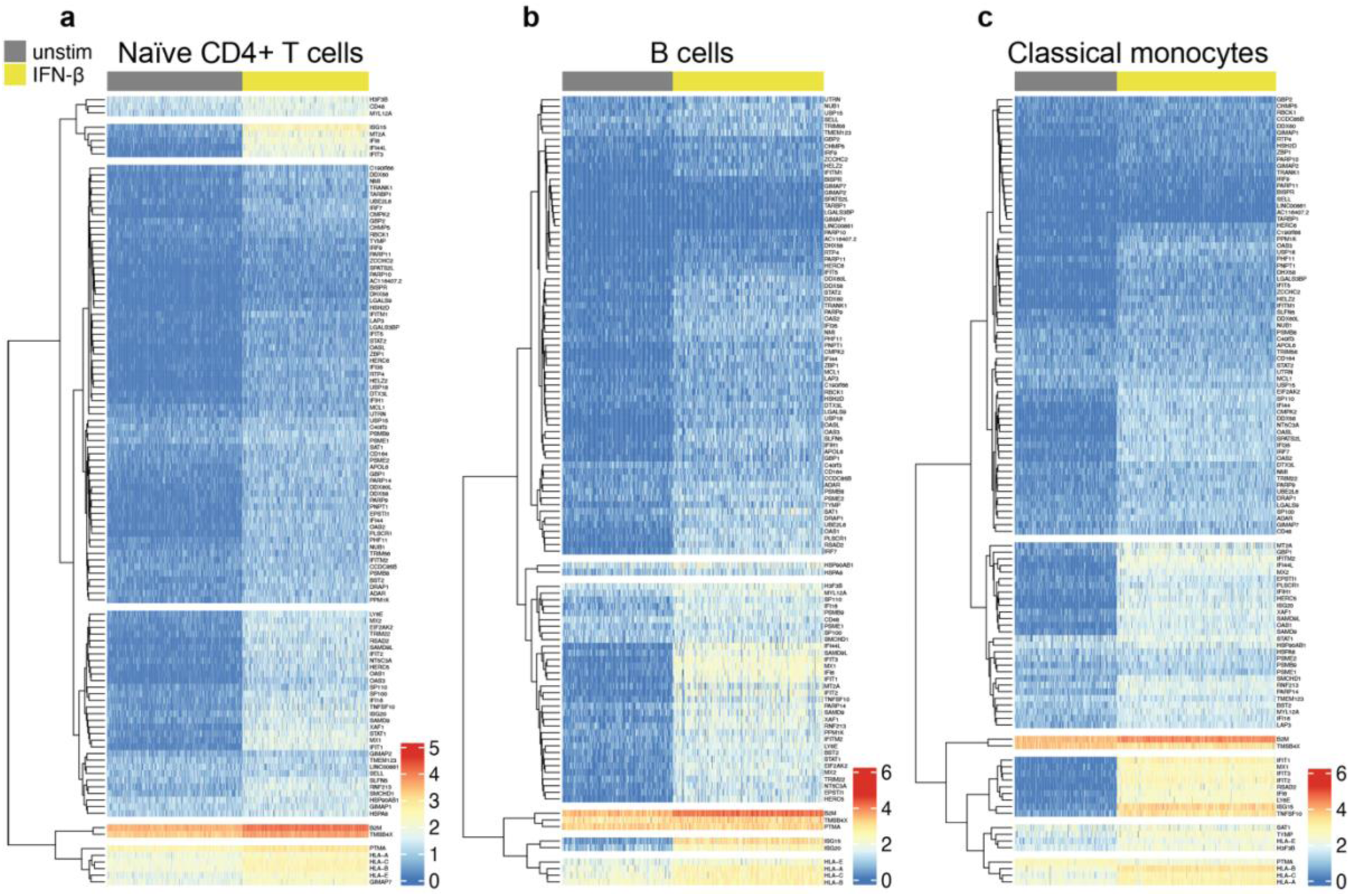
Response of naive CD4+ T cells to type I IFNs. (a-c) Heatmaps showing expression of the 113 genes significantly up-regulated by all tested type I IFNs in naïve CD4+ T cells, in unstimulated and IFN-β-treated cells for naïve CD4+ T cells (a), B cells (b) and classical monocytes (c). Gene names are provided for online viewing.

**Figure S24 (Related to Figure 6).**
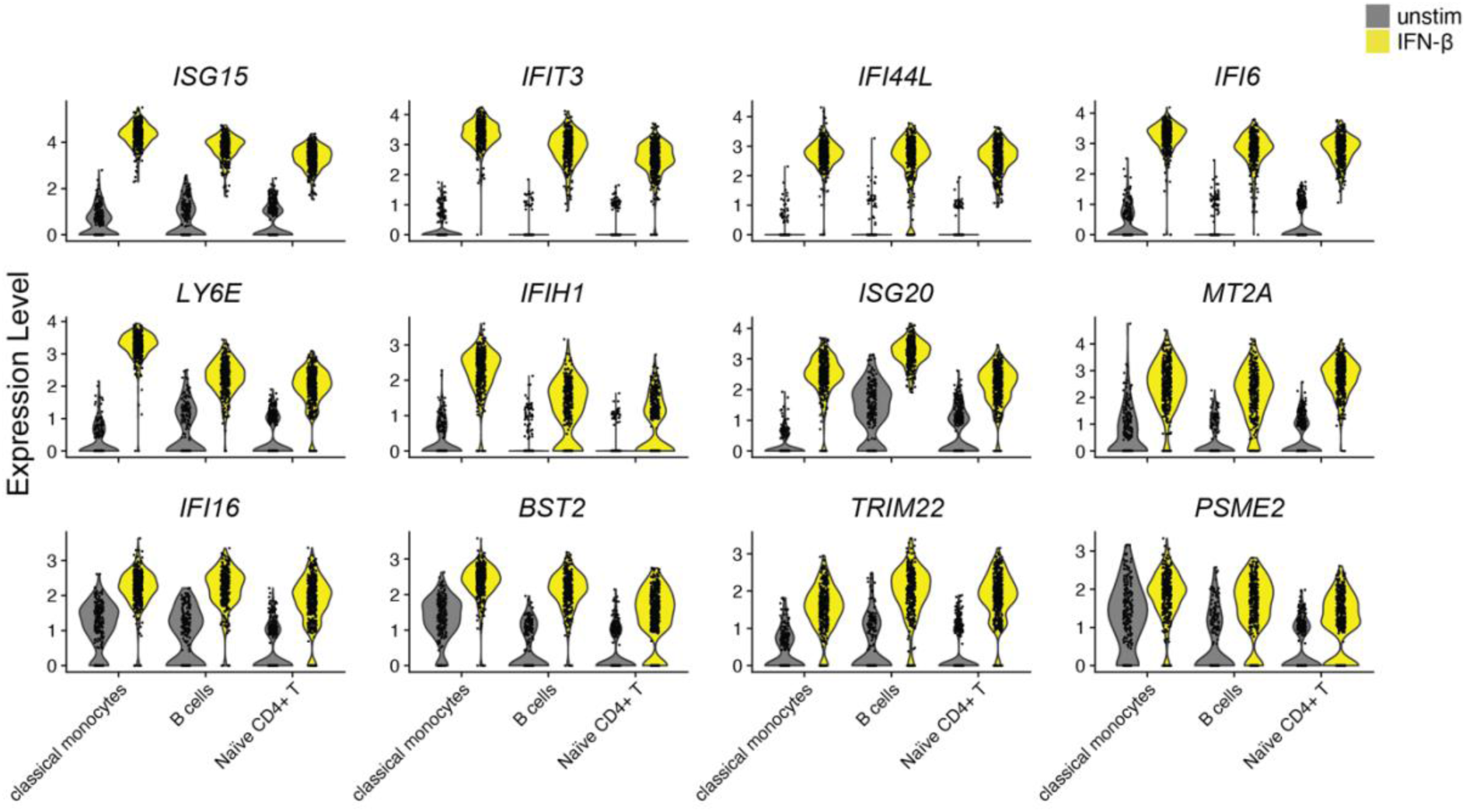
Selected ISGs significantly up-regulated by classical monocytes, B cells and naïve CD4+ T cells. Violin plots showing expression of twelve ISGs significantly up-regulated by classical monocytes, B cells and naïve CD4+ T cells in response to all type I IFN subtypes tested, in unstimulated and IFN-β-stimulated cells.

**Figure S25 (Related to Figure 6).**
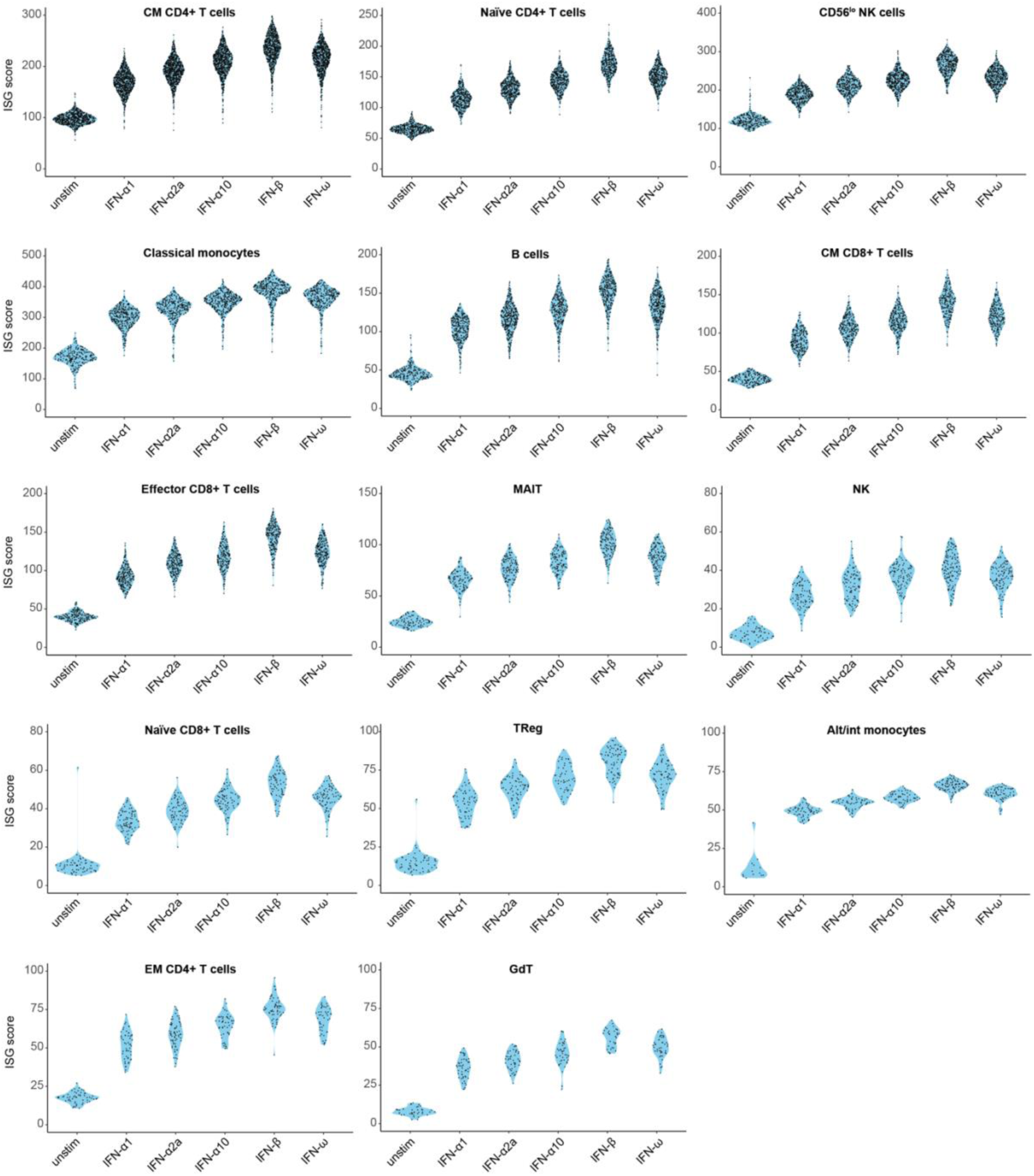
ISG scores for each cell type. Sinaplots with violin outline showing the ISG scores calculated using all genes significantly up-regulated by all type I IFNs for each cell type.

**Figure S26 (Related to Figure 6).**
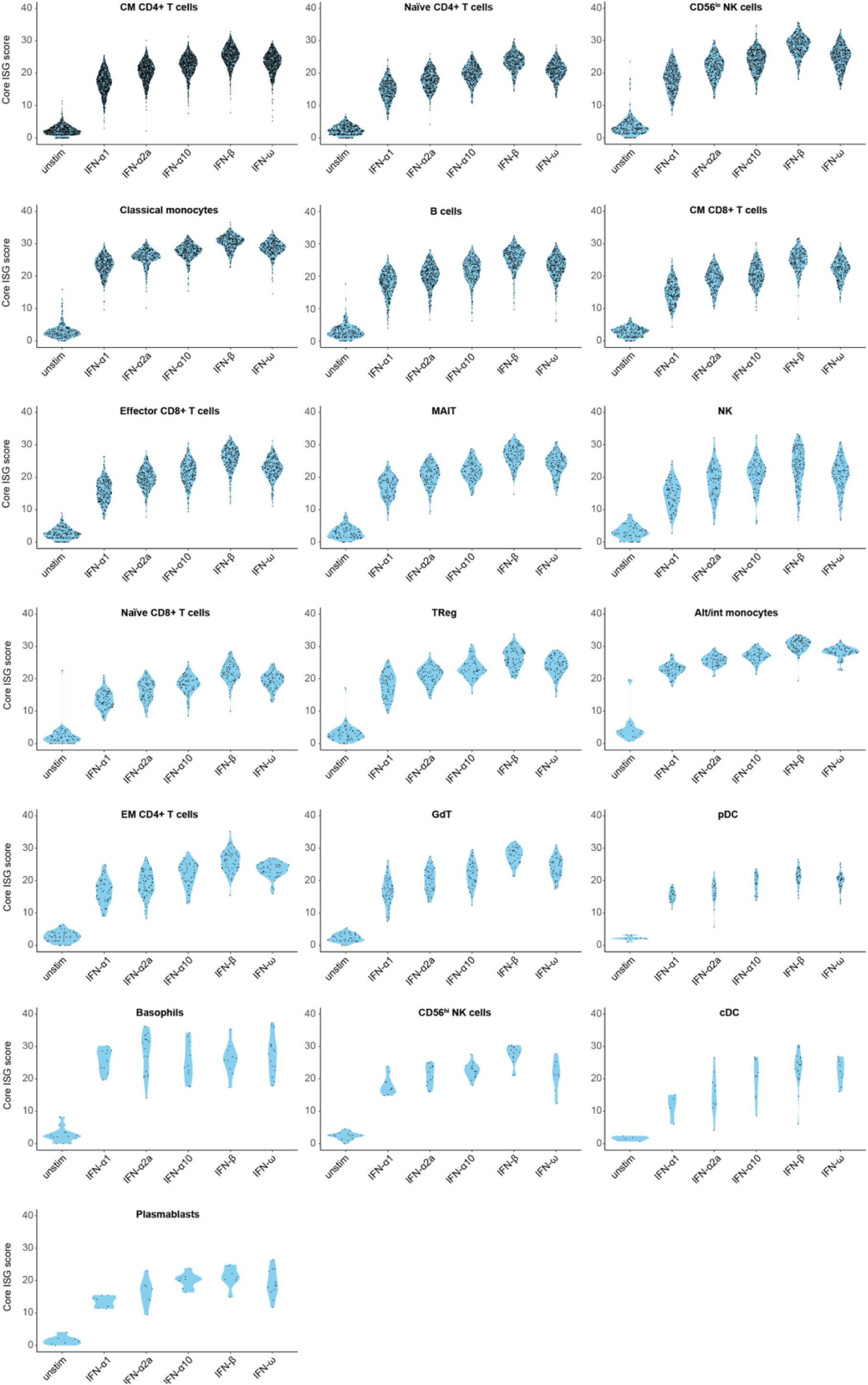
Core ISG scores for each cell type. Sinaplots with violin outline showing the ISG scores calculated using our ten core ISGs across each cell type.

## References

1. McNab, F., Mayer-Barber, K., Sher, A., Wack, A., and O’Garra, A. (2015). Type I interferons in infectious disease. Nature reviews. Immunology 15, 87–103. 10.1038/nri3787.

2. Crow, Y.J., and Manel, N. (2015). Aicardi-Goutieres syndrome and the type I interferonopathies. Nature reviews. Immunology 15, 429–440. 10.1038/nri3850.

3. Zitvogel, L., Galluzzi, L., Kepp, O., Smyth, M.J., and Kroemer, G. (2015). Type I interferons in anticancer immunity. Nature reviews. Immunology 15, 405–414. 10.1038/nri3845.

4. Isaacs, A., and Lindenmann, J. (1957). Virus interference. I. The interferon. Proc R Soc Lond B Biol Sci 147, 258–267.

5. Nagano, Y., and Kojima, Y. (1954). [Immunizing property of vaccinia virus inactivated by ultraviolets rays]. C R Seances Soc Biol Fil 148, 1700–1702.

6. Stark, G.R., and Darnell, J.E., Jr. (2012). The JAK-STAT pathway at twenty. Immunity 36, 503–514. 10.1016/j.immuni.2012.03.013.

7. Hartmann, G. (2017). Nucleic Acid Immunity. Adv Immunol 133, 121–169. 10.1016/bs.ai.2016.11.001.

8. Platanias, L.C. (2005). Mechanisms of type-I- and type-II-interferon-mediated signalling. Nature reviews. Immunology 5, 375–386. 10.1038/nri1604.

9. Yang, C.H., Shi, W., Basu, L., Murti, A., Constantinescu, S.N., Blatt, L., Croze, E., Mullersman, J.E., and Pfeffer, L.M. (1996). Direct association of STAT3 with the IFNAR-1 chain of the human type I interferon receptor. J Biol Chem 271, 8057–8061.

10. Schreiber, G. (2017). The molecular basis for differential type I interferon signaling. J Biol Chem 292, 7285–7294. 10.1074/jbc.R116.774562.

11. de Weerd, N.A., and Nguyen, T. (2012). The interferons and their receptors--distribution and regulation. Immunol Cell Biol 90, 483–491. 10.1038/icb.2012.9.

12. Schreiber, G., and Piehler, J. (2015). The molecular basis for functional plasticity in type I interferon signaling. Trends Immunol 36, 139–149. 10.1016/j.it.2015.01.002.

13. He, Y., Fu, W., Cao, K., He, Q., Ding, X., Chen, J., Zhu, L., Chen, T., Ding, L., Yang, Y., et al. (2020). IFN-kappa suppresses the replication of influenza A viruses through the IFNAR-MAPK-Fos-CHD6 axis. Sci Signal 13. 10.1126/scisignal.aaz3381.

14. Mazewski, C., Perez, R.E., Fish, E.N., and Platanias, L.C. (2020). Type I Interferon (IFN)-Regulated Activation of Canonical and Non-Canonical Signaling Pathways. Front Immunol 11, 606456. 10.3389/fimmu.2020.606456.

15. Rusinova, I., Forster, S., Yu, S., Kannan, A., Masse, M., Cumming, H., Chapman, R., and Hertzog, P.J. (2013). Interferome v2.0: an updated database of annotated interferon-regulated genes. Nucleic acids research 41, D1040–1046. 10.1093/nar/gks1215.

16. Schoggins, J.W., Wilson, S.J., Panis, M., Murphy, M.Y., Jones, C.T., Bieniasz, P., and Rice, C.M. (2011). A diverse range of gene products are effectors of the type I interferon antiviral response. Nature 472, 481–485. 10.1038/nature09907.

17. Shaw, A.E., Hughes, J., Gu, Q., Behdenna, A., Singer, J.B., Dennis, T., Orton, R.J., Varela, M., Gifford, R.J., Wilson, S.J., and Palmarini, M. (2017). Fundamental properties of the mammalian innate immune system revealed by multispecies comparison of type I interferon responses. PLoS Biol 15, e2004086. 10.1371/journal.pbio.2004086.

18. Guo, K., Shen, G., Kibbie, J., Gonzalez, T., Dillon, S.M., Smith, H.A., Cooper, E.H., Lavender, K., Hasenkrug, K.J., Sutter, K., et al. (2020). Qualitative Differences Between the IFNalpha subtypes and IFNbeta Influence Chronic Mucosal HIV-1 Pathogenesis. PLoS Pathog 16, e1008986. 10.1371/journal.ppat.1008986.

19. Malim, M.H., and Bieniasz, P.D. (2012). HIV Restriction Factors and Mechanisms of Evasion. Cold Spring Harb Perspect Med 2, a006940. 10.1101/cshperspect.a006940.

20. Schoggins, J.W. (2019). Interferon-Stimulated Genes: What Do They All Do? Annu Rev Virol 6, 567–584. 10.1146/annurev-virology-092818-015756.

21. Fung, K.Y., Mangan, N.E., Cumming, H., Horvat, J.C., Mayall, J.R., Stifter, S.A., De Weerd, N., Roisman, L.C., Rossjohn, J., Robertson, S.A., et al. (2013). Interferon-epsilon protects the female reproductive tract from viral and bacterial infection. Science 339, 1088–1092. 10.1126/science.1233321.

22. Barriga, F.M., Tsanov, K.M., Ho, Y.J., Sohail, N., Zhang, A., Baslan, T., Wuest, A.N., Del Priore, I., Meskauskaite, B., Livshits, G., et al. (2022). MACHETE identifies interferon-encompassing chromosome 9p21.3 deletions as mediators of immune evasion and metastasis. Nat Cancer 3, 1367–1385. 10.1038/s43018-022-00443-5.

23. LaFleur, D.W., Nardelli, B., Tsareva, T., Mather, D., Feng, P., Semenuk, M., Taylor, K., Buergin, M., Chinchilla, D., Roshke, V., et al. (2001). Interferon-kappa, a novel type I interferon expressed in human keratinocytes. J Biol Chem 276, 39765–39771. 10.1074/jbc.M102502200.

24. Wolf, S.J., Audu, C.O., Joshi, A., denDekker, A., Melvin, W.J., Davis, F.M., Xing, X., Wasikowski, R., Tsoi, L.C., Kunkel, S.L., et al. (2022). IFN-kappa is critical for normal wound repair and is decreased in diabetic wounds. JCI Insight 7. 10.1172/jci.insight.152765.

25. Pestka, S., Krause, C.D., and Walter, M.R. (2004). Interferons, interferon-like cytokines, and their receptors. Immunological reviews 202, 8–32. 10.1111/j.0105-2896.2004.00204.x.

26. Ortaldo, J.R., Herberman, R.B., Harvey, C., Osheroff, P., Pan, Y.C., Kelder, B., and Pestka, S. (1984). A species of human alpha interferon that lacks the ability to boost human natural killer activity. Proceedings of the National Academy of Sciences of the United States of America 81, 4926–4929.

27. Lavender, K.J., Gibbert, K., Peterson, K.E., Van Dis, E., Francois, S., Woods, T., Messer, R.J., Gawanbacht, A., Muller, J.A., Munch, J., et al. (2016). Interferon Alpha Subtype-Specific Suppression of HIV-1 Infection In Vivo. J Virol 90, 6001–6013. 10.1128/JVI.00451-16.

28. Harper, M.S., Guo, K., Gibbert, K., Lee, E.J., Dillon, S.M., Barrett, B.S., McCarter, M.D., Hasenkrug, K.J., Dittmer, U., Wilson, C.C., and Santiago, M.L. (2015). Interferon-alpha Subtypes in an Ex Vivo Model of Acute HIV-1 Infection: Expression, Potency and Effector Mechanisms. PLoS Pathog 11, e1005254. 10.1371/journal.ppat.1005254.

29. Tauzin, A., Espinosa Ortiz, A., Blake, O., Soundaramourty, C., Joly-Beauparlant, C., Nicolas, A., Droit, A., Dutrieux, J., Estaquier, J., and Mammano, F. (2021). Differential Inhibition of HIV Replication by the 12 Interferon Alpha Subtypes. J Virol 95, e0231120. 10.1128/JVI.02311-20.

30. Buzzai, A.C., Wagner, T., Audsley, K.M., Newnes, H.V., Barrett, L.W., Barnes, S., Wylie, B.C., Stone, S., McDonnell, A., Fear, V.S., et al. (2020). Diverse Anti-Tumor Immune Potential Driven by Individual IFNalpha Subtypes. Front Immunol 11, 542. 10.3389/fimmu.2020.00542.

31. Matos, A.D.R., Wunderlich, K., Schloer, S., Schughart, K., Geffers, R., Seders, M., Witt, M., Christersson, A., Wiewrodt, R., Wiebe, K., et al. (2019). Antiviral potential of human IFN-alpha subtypes against influenza A H3N2 infection in human lung explants reveals subtype-specific activities. Emerging microbes & infections 8, 1763–1776. 10.1080/22221751.2019.1698271.

32. Xie, X., Karakoese, Z., Ablikim, D., Ickler, J., Schuhenn, J., Zeng, X., Feng, X., Yang, X., Dittmer, U., Yang, D., et al. (2022). IFNalpha subtype-specific susceptibility of HBV in the course of chronic infection. Front Immunol 13, 1017753. 10.3389/fimmu.2022.1017753.

33. Schuhenn, J., Meister, T.L., Todt, D., Bracht, T., Schork, K., Billaud, J.N., Elsner, C., Heinen, N., Karakoese, Z., Haid, S., et al. (2022). Differential interferon-alpha subtype induced immune signatures are associated with suppression of SARS-CoV-2 infection. Proceedings of the National Academy of Sciences of the United States of America 119. 10.1073/pnas.2111600119.

34. Rout, S.S., Di, Y., Dittmer, U., Sutter, K., and Lavender, K.J. (2022). Distinct effects of treatment with two different interferon-alpha subtypes on HIV-1-associated T-cell activation and dysfunction in humanized mice. AIDS 36, 325–336. 10.1097/QAD.0000000000003111.

35. Karakoese, Z., Schwerdtfeger, M., Karsten, C.B., Esser, S., Dittmer, U., and Sutter, K. (2022). Distinct Type I Interferon Subtypes Differentially Stimulate T Cell Responses in HIV-1-Infected Individuals. Front Immunol 13, 936918. 10.3389/fimmu.2022.936918.

36. Schmitz, Y., Schwerdtfeger, M., Westmeier, J., Littwitz-Salomon, E., Alt, M., Brochhagen, L., Krawczyk, A., and Sutter, K. (2022). Superior antiviral activity of IFNbeta in genital HSV-1 infection. Front Cell Infect Microbiol 12, 949036. 10.3389/fcimb.2022.949036.

37. Thomas, C., Moraga, I., Levin, D., Krutzik, P.O., Podoplelova, Y., Trejo, A., Lee, C., Yarden, G., Vleck, S.E., Glenn, J.S., et al. (2011). Structural linkage between ligand discrimination and receptor activation by type I interferons. Cell 146, 621–632. 10.1016/j.cell.2011.06.048.

38. de Weerd, N.A., Vivian, J.P., Nguyen, T.K., Mangan, N.E., Gould, J.A., Braniff, S.J., Zaker-Tabrizi, L., Fung, K.Y., Forster, S.C., Beddoe, T., et al. (2013). Structural basis of a unique interferon-beta signaling axis mediated via the receptor IFNAR1. Nature immunology 14, 901–907. 10.1038/ni.2667.

39. Mostafavi, S., Yoshida, H., Moodley, D., LeBoite, H., Rothamel, K., Raj, T., Ye, C.J., Chevrier, N., Zhang, S.Y., Feng, T., et al. (2016). Parsing the Interferon Transcriptional Network and Its Disease Associations. Cell 164, 564–578. 10.1016/j.cell.2015.12.032.

40. Uhlen, M., Karlsson, M.J., Zhong, W., Tebani, A., Pou, C., Mikes, J., Lakshmikanth, T., Forsstrom, B., Edfors, F., Odeberg, J., et al. (2019). A genome-wide transcriptomic analysis of protein-coding genes in human blood cells. Science 366. 10.1126/science.aax9198.

41. Derrien, T., Johnson, R., Bussotti, G., Tanzer, A., Djebali, S., Tilgner, H., Guernec, G., Martin, D., Merkel, A., Knowles, D.G., et al. (2012). The GENCODE v7 catalog of human long noncoding RNAs: analysis of their gene structure, evolution, and expression. Genome Res 22, 1775–1789. 10.1101/gr.132159.111.

42. Zheng, H., Brennan, K., Hernaez, M., and Gevaert, O. (2019). Benchmark of long non-coding RNA quantification for RNA sequencing of cancer samples. Gigascience 8. 10.1093/gigascience/giz145.

43. Suarez, B., Prats-Mari, L., Unfried, J.P., and Fortes, P. (2020). LncRNAs in the Type I Interferon Antiviral Response. Int J Mol Sci 21. 10.3390/ijms21176447.

44. Stoeckius, M., Hafemeister, C., Stephenson, W., Houck-Loomis, B., Chattopadhyay, P.K., Swerdlow, H., Satija, R., and Smibert, P. (2017). Simultaneous epitope and transcriptome measurement in single cells. Nature methods 14, 865–868. 10.1038/nmeth.4380.

45. Hao, Y., Hao, S., Andersen-Nissen, E., Mauck, W.M., 3rd, Zheng, S., Butler, A., Lee, M.J., Wilk, A.J., Darby, C., Zager, M., et al. (2021). Integrated analysis of multimodal single-cell data. Cell 184, 3573–3587 e3529. 10.1016/j.cell.2021.04.048.

46. Rue-Albrecht, K., Marini, F., Soneson, C., and Lun, A. (2018). iSEE: Interactive SummarizedExperiment Explorer [version 1; peer review: 3 approved]. F1000Research 7. 10.12688/f1000research.14966.1.

47. Mattick, J.S., and Rinn, J.L. (2015). Discovery and annotation of long noncoding RNAs. Nat Struct Mol Biol 22, 5–7. 10.1038/nsmb.2942.

48. Statello, L., Guo, C.J., Chen, L.L., and Huarte, M. (2021). Gene regulation by long non-coding RNAs and its biological functions. Nat Rev Mol Cell Biol 22, 96–118. 10.1038/s41580-020-00315-9.

49. Ji, X., Meng, W., Liu, Z., and Mu, X. (2022). Emerging Roles of lncRNAs Regulating RNA-Mediated Type-I Interferon Signaling Pathway. Front Immunol 13, 811122. 10.3389/fimmu.2022.811122.

50. Kambara, H., Niazi, F., Kostadinova, L., Moonka, D.K., Siegel, C.T., Post, A.B., Carnero, E., Barriocanal, M., Fortes, P., Anthony, D.D., and Valadkhan, S. (2014). Negative regulation of the interferon response by an interferon-induced long non-coding RNA. Nucleic Acids Res 42, 10668–10680. 10.1093/nar/gku713.

51. Mariotti, B., Servaas, N.H., Rossato, M., Tamassia, N., Cassatella, M.A., Cossu, M., Beretta, L., van der Kroef, M., Radstake, T., and Bazzoni, F. (2019). The Long Non-coding RNA NRIR Drives IFN-Response in Monocytes: Implication for Systemic Sclerosis. Front Immunol 10, 100. 10.3389/fimmu.2019.00100.

52. Song, L., Chen, J., Lo, C.Z., Guo, Q., Consortium, Z.I.B., Feng, J., and Zhao, X.M. (2022). Impaired type I interferon signaling activity implicated in the peripheral blood transcriptome of preclinical Alzheimer’s disease. EBioMedicine 82, 104175. 10.1016/j.ebiom.2022.104175.

53. Bayyurt, B., Bakir, M., Engin, A., Oksuz, C., and Arslan, S. (2021). Investigation of NEAT1, IFNG-AS1, and NRIR expression in Crimean-Congo hemorrhagic fever. J Med Virol 93, 3300-3304. 10.1002/jmv.26606.

54. Pavlovich, S.S., Lovett, S.P., Koroleva, G., Guito, J.C., Arnold, C.E., Nagle, E.R., Kulcsar, K., Lee, A., Thibaud-Nissen, F., Hume, A.J., et al. (2018). The Egyptian Rousette Genome Reveals Unexpected Features of Bat Antiviral Immunity. Cell 173, 1098–1110 e1018. 10.1016/j.cell.2018.03.070.

55. Zhou, P., Tachedjian, M., Wynne, J.W., Boyd, V., Cui, J., Smith, I., Cowled, C., Ng, J.H., Mok, L., Michalski, W.P., et al. (2016). Contraction of the type I IFN locus and unusual constitutive expression of IFN-alpha in bats. Proceedings of the National Academy of Sciences of the United States of America 113, 2696–2701. 10.1073/pnas.1518240113.

56. Hardy, M.P., Owczarek, C.M., Jermiin, L.S., Ejdeback, M., and Hertzog, P.J. (2004). Characterization of the type I interferon locus and identification of novel genes. Genomics 84, 331–345. 10.1016/j.ygeno.2004.03.003.

57. Owczarek, C.M., Hwang, S.Y., Holland, K.A., Gulluyan, L.M., Tavaria, M., Weaver, B., Reich, N.C., Kola, I., and Hertzog, P.J. (1997). Cloning and characterization of soluble and transmembrane isoforms of a novel component of the murine type I interferon receptor, IFNAR 2. J Biol Chem 272, 23865–23870. 10.1074/jbc.272.38.23865.

58. Symons, J.A., Alcami, A., and Smith, G.L. (1995). Vaccinia virus encodes a soluble type I interferon receptor of novel structure and broad species specificity. Cell 81, 551–560. 10.1016/0092-8674(95)90076-4.

59. Honda, K., Takaoka, A., and Taniguchi, T. (2006). Type I interferon [corrected] gene induction by the interferon regulatory factor family of transcription factors. Immunity 25, 349–360. 10.1016/j.immuni.2006.08.009.

60. Marie, I., Durbin, J.E., and Levy, D.E. (1998). Differential viral induction of distinct interferon-alpha genes by positive feedback through interferon regulatory factor-7. The EMBO journal 17, 6660–6669. 10.1093/emboj/17.22.6660.

61. Izaguirre, A., Barnes, B.J., Amrute, S., Yeow, W.S., Megjugorac, N., Dai, J., Feng, D., Chung, E., Pitha, P.M., and Fitzgerald-Bocarsly, P. (2003). Comparative analysis of IRF and IFN-alpha expression in human plasmacytoid and monocyte-derived dendritic cells. J Leukoc Biol 74, 1125–1138. 10.1189/jlb.0603255.

62. Kerkmann, M., Rothenfusser, S., Hornung, V., Towarowski, A., Wagner, M., Sarris, A., Giese, T., Endres, S., and Hartmann, G. (2003). Activation with CpG-A and CpG-B oligonucleotides reveals two distinct regulatory pathways of type I IFN synthesis in human plasmacytoid dendritic cells. J Immunol 170, 4465–4474. 10.4049/jimmunol.170.9.4465.

63. Nienaltowski, K., Rigby, R.E., Walczak, J., Zakrzewska, K.E., Glow, E., Rehwinkel, J., and Komorowski, M. (2021). Fractional response analysis reveals logarithmic cytokine responses in cellular populations. Nature communications 12, 4175. 10.1038/s41467-021-24449-2.

64. Lamble, S., Batty, E., Attar, M., Buck, D., Bowden, R., Lunter, G., Crook, D., El-Fahmawi, B., and Piazza, P. (2013). Improved workflows for high throughput library preparation using the transposome-based Nextera system. BMC Biotechnol 13, 104. 10.1186/1472-6750-13-104.

65. Stoeckius, M., Zheng, S., Houck-Loomis, B., Hao, S., Yeung, B.Z., Mauck, W.M., 3rd, Smibert, P., and Satija, R. (2018). Cell Hashing with barcoded antibodies enables multiplexing and doublet detection for single cell genomics. Genome Biol 19, 224. 10.1186/s13059-018-1603-1.

66. Chevrier, S., Crowell, H.L., Zanotelli, V.R.T., Engler, S., Robinson, M.D., and Bodenmiller, B. (2018). Compensation of Signal Spillover in Suspension and Imaging Mass Cytometry. Cell Syst 6, 612–620 e615. 10.1016/j.cels.2018.02.010.

67. Nowicka, M., Krieg, C., Crowell, H.L., Weber, L.M., Hartmann, F.J., Guglietta, S., Becher, B., Levesque, M.P., and Robinson, M.D. (2017). CyTOF workflow: differential discovery in high-throughput high-dimensional cytometry datasets. F1000Res 6, 748. 10.12688/f1000research.11622.3.

68. Cribbs, A., Luna-Valero, S., George, C., Sudbery, I., Berlanga-Taylor, A., Sansom, S., Smith, T., Ilott, N., Johnson, J., Scaber, J., et al. (2019). CGAT-core: a python framework for building scalable, reproducible computational biology workflows [version 2; peer review: 1 approved, 1 approved with reservations]. F1000Research 8. 10.12688/f1000research.18674.2.

69. Volders, P.J., Anckaert, J., Verheggen, K., Nuytens, J., Martens, L., Mestdagh, P., and Vandesompele, J. (2019). LNCipedia 5: towards a reference set of human long non-coding RNAs. Nucleic acids research 47, D135–D139. 10.1093/nar/gky1031.

70. Soneson, C., Love, M.I., and Robinson, M.D. (2015). Differential analyses for RNA-seq: transcript-level estimates improve gene-level inferences. F1000Res 4, 1521. 10.12688/f1000research.7563.2.

71. Love, M.I., Huber, W., and Anders, S. (2014). Moderated estimation of fold change and dispersion for RNA-seq data with DESeq2. Genome Biol 15, 550. 10.1186/s13059-014-0550-8.

72. Young, M.D., Wakefield, M.J., Smyth, G.K., and Oshlack, A. (2010). Gene ontology analysis for RNA-seq: accounting for selection bias. Genome Biol 11, R14. 10.1186/gb-2010-11-2-r14.

73. Roelli, P., bbimber, Flynn, B., santiagorevale, and Gui, G. (2019). Hoohm/CITE-seq-Count: 1.4.2. Zenodo. https://doi.org/10.5281/zenodo.2590196.

